# High-resolution cryo-EM structures of a protein pore reveal diverse roles of membrane lipids

**DOI:** 10.1101/2024.06.26.600493

**Authors:** Gašper Šolinc, Marija Srnko, Franci Merzel, Ana Crnković, Mirijam Kozorog, Marjetka Podobnik, Gregor Anderluh

## Abstract

The structure and function of membrane proteins depend on their interactions with the lipid molecules that constitute lipid membranes. Actinoporins are a family of α-pore-forming proteins that bind specifically to sphingomyelin-containing lipid membranes, where they oligomerize and form transmembrane pores. The numerous contacts they form with the lipid membrane make them an exemplary object for studying the different roles that lipids play in the structure and function of membrane proteins. Through a comprehensive cryo-electron microscopic analysis of a pore formed by an actinoporin Fav from the coral *Orbicella faveolata*, we show that the octameric pore interacts with 112 lipids in the upper leaflet of the membrane. The structures of Fav pores formed on different lipid membranes reveal the different roles of lipids and demonstrate that the actinoporin surface is perfectly suited for binding multiple receptor sphingomyelin molecules. When cholesterol is present in the membrane, it forms nanodomains associated with the pore, leading to a tighter arrangement of lipids, which in turn increases the stability of the pores. Atomistic simulations support the structural data, show that the protein-bound lipids are not mobile, and reveal additional effects of the pore on the lipid membrane. Overall, these data reveal a complex network of protein-lipid and lipid-lipid interactions, and an underrated role of lipids in the structure and function of transmembrane protein complexes.

## Main text

Lipid membranes are crucial for the functioning of organisms. A significant portion of the mammalian proteome is associated with lipid membranes^1^ and many membrane-associated proteins are valuable drug targets^2^. It is becoming increasingly clear that lipids can affect proteins in many different ways, particularly in regulating the structure and function of membrane proteins^3-7^. For example, membrane lipids can also act as cofactors required for the proper functioning of membrane enzymes^8^ or as structural support in the formation of membrane protein assemblies^9^. Lipids affect transporters^10,11^ or ion channel gating and activity by directly interacting with the protein or indirectly by altering the properties of the membrane^12,13^. They play an important role in regulating the activity of G-protein-coupled receptors, such as serotonin receptors^14^ and β2-adrenergic receptors^15^, through specific binding of phospholipids or through cholesterol molecules surrounding the receptor’s transmembrane domain influencing the ligand-binding pocket.

Due to the nature of protein-lipid interactions and the dynamics of lipid molecules, the determination of protein-lipid interactions at high resolution is a major challenge^16,17^. Thanks to advances in cryo-electron microscopy (cryo-EM), the number of known structures of membrane proteins with associated lipids continues to increase^17,18^. The lipids observed in these structures are usually one of two types, a single, specifically bound lipid with a well-defined density or multiple ordered acyl chains without defined headgroups in a cleft or at the interface between subunits of a larger complex^19,20^. In several cases lipids have been unambiguously identified in three dimensional structures of membrane protein complexes, providing valuable information about their effects on protein structure and function^14,21-23^.

The crystal structure of the octameric transmembrane pore of the pore-forming toxin, an actinoporin fragaceatoxin C (FraC)^24^ shows the positions of the headgroups of three lipids bound to a single protomer. However, how the membrane is organized around the pore and how the pore itself influences the arrangement of the membrane lipids has not yet been shown in detail. Here, we used a homologue of FraC from the coral *Orbicella faveolata*, Fav, to specifically identify and reconstruct by cryo-EM the atomic model of lipid molecules associated with the transmembrane pore. We developed a protocol to produce stable soluble pores retaining a significant number of bound lipids, which we were able to structurally resolve and assign to different functional roles. In combination with molecular dynamics (MD) simulations, we uncovered extensive lipid-protein and lipid-lipid interactions, and gained unprecedented insight into the embedding of the protein complex in the lipid membrane.

## Preparation of soluble Fav pores

Fav is a homologue of actinoporins with unique extensions at the N- and C-termini (Extended Data Fig. 1a). We determined the crystal structure of monomeric Fav with truncated 53 N-terminal residues (ΔN53Fav), as this region was predicted by Swiss Model^25^ to be unstructured.

The electron density map was defined only for the residues beyond A76. The structured part of ΔN53Fav showed a clear conservation of a typical actinoporin monomer consisting of a central β-sandwich flanked by two α-helices (Extended Data Fig. 1b and c, Extended Data Table 1)^26^. The C-terminal extension (E254-M259) was completely resolved and anchored to the β-sandwich by a disulfide bond, unique among actinoporins (Extended Data Fig. 1b).

For cryo-EM analysis, Fav pores were prepared by incubating monomeric wild-type Fav with 1,2-dioleoyl-*sn*-glycero-3-phosphocholine (DOPC):sphingomyelin (SM) 1:1 (mol:mol) large unilamellar vesicles (LUVs) (Extended Data Fig. 1d). We observed pores of clearly distinguishable octameric and nonameric stoichiometries (Extended Data Fig. 1d). We tested different detergents to solubilize the pores and finally solubilized them with lauryl dimethylamine oxide. We separated the solubilized pores from excess lipids and detergents by ion-exchange chromatography and concentrated them before further use (Extended Data Fig. 1e).

## The cryo-EM structure of solubilized Fav pores reveals a layer of lipids supporting the pore structure

3D cryo-EM reconstruction was successful only for octameric pores, indicating low stability of the isolated nonameric pores (Fig. 1a, Extended Data Fig. 2). The funnel-shaped transmembrane channel, built of a cluster of eight amphipathic α1-helices (residues A80-N103) that detach from the central β-sandwich cores of protomers during pore formation^27^, is embedded in a micelle composed of detergents and remnant lipids. The residues upstream the N-terminal α1-helix (G1-I78) are not defined by cryo-EM density. While the overall architecture of the Fav pore determined at 2.6 Å resolution, including the cap region and the transmembrane helical cluster (Fig. 1b), is similar to that of FraC^24^, there are the following notable differences. In contrast to the FraC pore, the electrostatic potential of the transmembrane channel surface of the Fav pore is predominantly negative (Extended Data Fig. 3a). The transmembrane channel of the Fav pore is wider than that of the FraC pore, with the narrowest part of 1.54 nm in diameter located at the constriction of the channel formed by the cluster of E88 side chains (Extended Data Fig. 3a). Finally, the elongation at the C-terminus of Fav provides additional contacts between the protomers (Extended Data Fig. 3a).

**Fig. 1.**
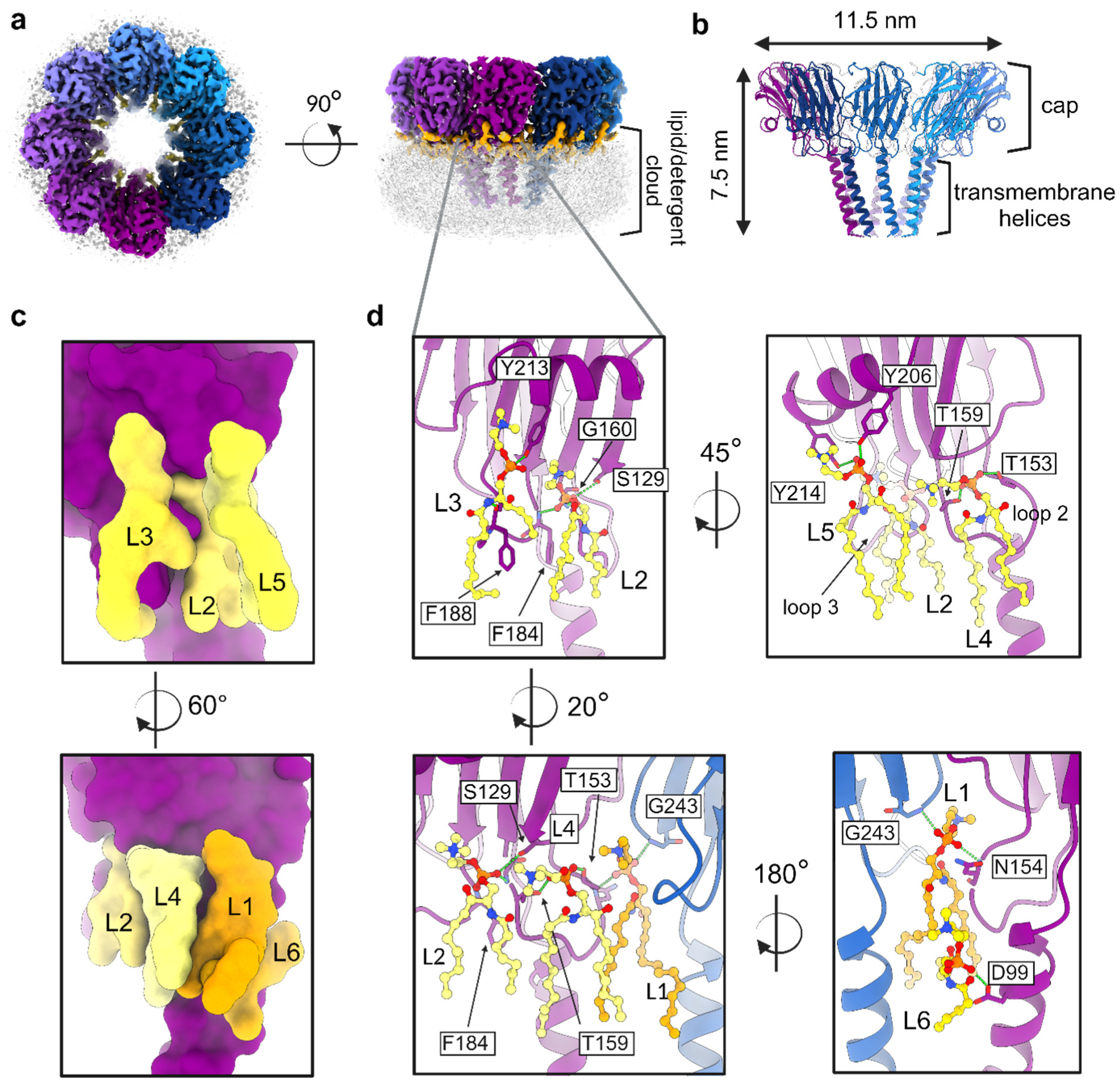
The structure of the Fav pore formed on DOPC:SM (1:1) membranes. **a**, A top and a side view of cryo-EM map of octameric Fav pore extracted from DOPC:SM large unilamellar vesicles. Each protomer is coloured with a distinct colour, regions of the map corresponding to well-defined lipids are shown in orange. The transparent grey cloud around the bottom part of the pore is a cryo-EM density of the lipid/detergent micelle. **b**, Cartoon representation of the Fav pore. The cap region and transmembrane (TM) helical cluster and pore dimensions are marked. **c**, Surface representation of a single Fav pore protomer with marked six lipid positions (L1-L6; orange and yellow) identified from the pore structure (purple). **d**, Detailed look at the lipid positions and their coordination.

Three lipid molecules per protomer with partially resolved acyl chains were previously observed in the FraC pore (Extended Data Fig. 3b and c)^24^. In the Fav pore, we clearly observed lipid densities at the corresponding positions, labelled L1-L3 (Fig. 1c and d, Extended Data Fig. 3b and c). L1 is located between two neighbouring protomers. The polar headgroup of L1 is almost completely buried between the protomers in the pore, with its phosphate group forming a hydrogen bond with the backbone amide nitrogen and carbonyl oxygen of N154 of one protomer and the backbone amide nitrogen of G243 of the neighbouring protomer (Extended Data Fig. 4). The choline headgroup of SM forms cation-π interaction with Y186^28^. The acyl chains of L1 are additionally stabilized by hydrophobic interactions with the acyl chains of L4 and L6. L2 and L3 bind to a single protomer. L2 is located below the C-terminal α2-helix and between loops 2 and 3, where its phosphate group forms a hydrogen bond with the side chain of S129 and the backbone amide of G160 and F184. Its choline group forms cation-π interactions with Y189 and Y213. L2 also interacts with lipids L5, and L1 of the neighbouring protomer. The headgroup of L3 located in the pocket below the α2-helix and the acyl chains is supported by the hydrophobic tip of the Fav loop 3 (Fig. 1d, Extended Data Fig. 4). It forms hydrogen bonds with backbone carboxyl group of F188 and side chain of Y213 and cation-π interaction with Y213. The acyl chains of L3 are stabilized by hydrophobic interactions with the protein.

In addition to L1-L3, we observed densities corresponding to three other lipids in the Fav pore, L4-L6. L4 is located near loop 2 at the interface between two protomers, and is flanked by L1 and L2. The phosphate group of L4 forms two hydrogen bonds with the side chains of T153 and T159 and cation-π interaction with F128. L5 is located at the outer edge of the pore below the α2-helix, almost completely covering L2 (Fig. 1d). Its phosphate group forms hydrogen bonds with the side chains of Y206 and Y214 from the α2-helix and also cation-π interaction with Y214 (Extended Data Fig. 4). L6 is buried between the neighbouring protomers, with its headgroup pointing towards the pore lumen through the fenestration. Its phosphate group forms a hydrogen bond with the side chain of D99, and the acyl chains are stabilized by hydrophobic interactions with adjacent transmembrane α1-helices and L1 (Fig. 1d, Extended Data Fig. 4). L1 and L6 are buried into the pore between the protomers and are therefore referred to as structural lipids (Extended Data Fig. 5). L2 and L3 are located at positions previously proposed as important for actinoporin membrane binding^24^ and, together with L4 and L5, are thus referred to as receptor lipids. The headgroups of L1-L5 are well defined (Extended Data Fig. 4) and correspond to the phosphocholine headgroups of DOPC or SM used in vesicles. The densities corresponding to lipid tails can fit acyl chains of 3 (L3) to 12 carbon atoms (L1). The region under the cap of the pore contained no density (Extended Data Fig. 6a). In this void, we could observe alternative conformations of the acyl chains of L2 and L4 (Extended Data Fig. 6b and c).

Overall, the isolated wild-type pore of Fav is a protein-lipid pore complex consisting of 8 homoprotomers to which 48 lipids are stably bound. The lipid monolayer is bound to the protein by hydrogen bonds, cation-π interactions, and hydrophobic interactions, which are additionally supported by the hydrophobic interactions between the lipid tails.

## Cholesterol forms ordered nanodomains under the cap of the Fav pore

The activity of actinoporins is enhanced in the presence of cholesterol^29^. Therefore, our next goal was to investigate how cholesterol affects lipid arrangement and the pore structure. Furthermore, we could not resolve which lipid is bound in the pores formed on DOPC:SM membranes, as the DOPC and SM headgroups have the same phosphocholine structure. For this reason, we used 1-palmitoyl-2-oleoyl-*sn*-glycero-3-phospho-(1’-rac-glycerol) (POPG) instead of DOPC, which has a different head group than SM and should allow differentiation of the bound lipids. We prepared wild-type Fav pores on liposomes composed of POPG:SM:cholesterol 1:1:1 (mol:mol:mol). Cryo-EM reconstruction at 2.9 Å resolution showed a similar architecture of the pore as found in DOPC:SM membranes (Fig. 2a, Extended Data Fig. 7). However, the lipid region below the pore cap was significantly larger and better defined (Extended Data Fig. 8). We were able to resolve 15 lipids associated with each protomer, including 4 cholesterol molecules (CH1-CH4), 10 phospholipids (L1-L10) and one acyl chain (L11) per protomer.

**Fig. 2.**
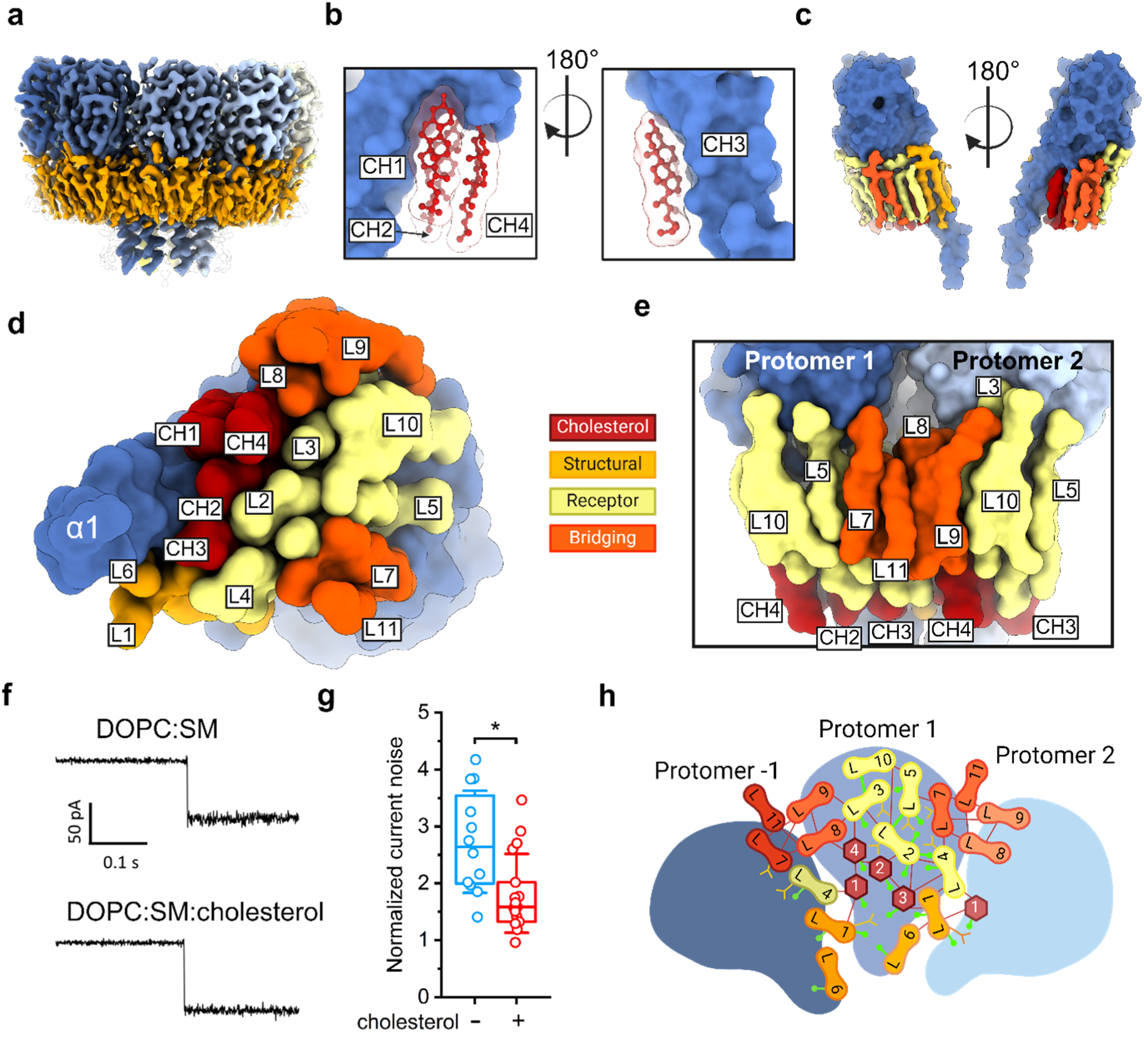
The structure of Fav pore prepared on POPG:SM:cholesterol (1:1:1) large unilamellar vesicles. **a**, A side view of cryo-EM map of octameric Fav pore. Parts of the map corresponding to individual pore protomers are coloured differently, and regions corresponding to well-defined lipids are shown in orange. **b**, Stick and ball representation of the four cholesterol (CH) molecules in red, overlaid by the corresponding molecular surface. Pore surface is in blue. **c** Surface representation of a Fav pore promoter with parts of the map corresponding to lipids associated with it shown at two different angles. The lipids are coloured according to the assigned function as shown in **d**. **d**, A view of a Fav pore protomer in the surface presentation from the bottom of an α1-helix, showing the positions of lipids defined in the cryo-EM structure of the pore. Lipids are coloured according to the assigned function. **e,** Surface representation of phospholipids resolved in the pore structure, highlighting the bridging lipids. **f**, Comparison of discrete ion current steps, corresponding to pore insertion induced by the addition of a monomeric Fav protein at a voltage of - 50 mV in two different lipid membrane bilayers as indicated. **g**, Normalized current noise for membranes from **f** in the absence (-) or presence (+) of cholesterol. n=12 and 17 single pore measurements, respectively. Two-sample t-test was used for statistical analysis. * p<0.01. **h**, A schematic representation of lipids and proteins observed in cryo-EM structure of the Fav pore extracted from POPG:SM:CH membrane. Hydrogen bonds between lipids are denoted by green lines, hydrogen bonds between lipids and proteins are denoted by a line with the circle at one end. Hydrophobic interactions between lipids are shown as red lines. Cation-π interactions between lipids and protein are indicated by orange inverted arrows.

A unique feature of the pores extracted from POPG:SM:cholesterol membranes was a patch of four cholesterol molecules (CH1-CH4) located between the transmembrane α1-helix of each protomer and the membrane binding loops 2 and 3 (Fig. 2b), and surrounded by lipids L1-L4 and L8. In all four cholesterol molecules, the hydroxyl groups point towards the bottom of the pore cap and form hydrogen bonds with the protomer (Extended Data Fig. 9). The ordered cholesterol nanodomain with a van der Waals volume of 1311 Å^3^ fills the space under the cap of the pore, resulting in a more ordered arrangement of the acyl chains of the phospholipids bound to the pore compared to the pores from DOPC:SM membranes. Consequently, in contrast to the pores from DOPC:SM membranes, L2 and L4 adopt a single conformation.

In the presence of cholesterol, L1-L5 adopt similar positions as in the pores extracted from DOPC:SM membranes. L6 appears to be shifted up for about 8 Å along the axis of the α1-helix, which is due to a different conformation of one of the acyl chains of the neighbouring L1 acyl group in the presence of cholesterol (Extended Data Fig. 10). Moreover, we were able to build four additional phospholipids (L7-L10) per protomer and an additional acyl chain (L11) (Fig 2c and d, Extended Data Fig. 11). L7-L9 are located between two protomers (Fig. 2d and e), however, with very limited contacts with them. L7 is located on one side of the protomer, and L8 and L9 are on the other side (Fig. 2d and e). The choline headgroup of L7 points towards the side chain of Y206 and is stabilised via cation-π interactions^28^, and additionally stabilised by hydrophobic interactions with the surrounding lipids L4, L5 and L11 of the same protomer and to some extent also with L8 and L9 of the neighbouring protomer (Fig. 2d and e, Extended Data Fig. 11). The headgroup of L8 is supported by the residues of the loop 3 and its acyl chains by CH4, L3 and L9 assigned to the same protomer, and L4 and L7 from the neighbouring protomers (Extended Data Fig. 11). L9 is located in the most outer lipid shell with almost no interaction surface with the protein, and is stabilised by lipid-lipid interactions with L3, L8 and L10 assigned to the same protomer and L7 and L11 from the neighbouring protomer (Extended Data Fig. 11, Extended Data Table 2). L10 is also located in the outer shell of the lipid binding region, between L5 and L9 (Fig. 2d). L10 choline headgroup is nestled between choline headgroup of L3 and side chain of Y214 from α2-helix, and its acyl chain is stabilized by extensive interactions with the acyl chains of L3 (Extended Data Fig. 11, Extended Data Table 2). L11 could only be modelled as an acyl chain at the outer lipid shell (Fig. 2d and e), bridging L7 of the same protomer with L8 and L9 of the neighbouring protomer (Extended Data Fig. 11). We have termed L7-L9 and L11 as bridging lipids because they have very little or no contact with the protomers. However, these lipids are in extensive contact with each other, and thus indirectly reinforce the pore structure via a protomer-1-lipid-lipid-protomer-2 bridge (Fig. 2e).

The presence of cholesterol in the membrane only slightly affects the position of the transmembrane helices (Extended Data Fig. 12). However, it has a small but significant effect on the ionic currents flowing through the pore. The increase in current noise upon pore formation was lower in cholesterol-containing membranes compared to cholesterol-free membranes (Fig. 2f and g).

The pores prepared with POPG:SM:cholesterol membranes, in combination with the good local resolution of the cryo-EM map, allowed us to identify lipids L1-L5 as SM. The map quality of the headgroup region of L6-L11 is not high enough to distinguish between POPG and SM headgroups. In the presence of cholesterol, the acyl chains of the lipids are better defined and are now for most of the observed lipids longer than 10 carbon atoms, reaching 15 carbon atoms in L1, L2 and L5. A large part of the protomer surface is involved in membrane interaction. Of the total available 9,339 Å^2^ surface area of the single protomer, the amphipathic transmembrane α1-helix occupies about 2,000 Å^2^. Of the remaining surface area, 1,500 Å^2^ is involved in protomer-protomer interactions, and 2,106 Å^2^ in protein-lipid headgroup interactions (Extended Data Table 2). Thus, a considerable part of the protomer surface is suitable for lipid binding and is used for this purpose. Overall, we were able resolve 15 lipids associated with each protomer in the Fav pore. This monolayer patch under the pore cap is fixed by a complex network of hydrogen bonds, hydrophobic interactions and cation-π interactions. Fifteen hydrogen bonds are formed between protein and lipids and two between lipids (L2-L5 and L3-L10) (Fig. 2h).

To confirm that cholesterol and not POPG is responsible for the tighter lipid arrangement, we solved the structure of the octameric pore extracted from DOPC:SM:cholesterol 1:1:1 (mol:mol:mol) LUVs at 3.1 Å resolution (Extended Data Fig. 13a). The overall structure of the pore is almost identical with 9 phospholipids and 4 cholesterol molecules visible at comparable positions as in the pores extracted from POPG:SM:cholesterol vesicles (Extended Data Fig. 13b and c). In addition, we prepared pores on lipid nanodiscs to rule out the possibility that the observed lipid positions around the pore are an artefact of membrane solubilization with detergents. Nanodiscs composed of DOPC:SM:cholesterol 1:1:1 (mol:mol:mol) were incubated with monomeric wild-type Fav and then analysed by cryo-EM. We were able to reconstruct the octameric pore structure with an overall resolution of 3.6 Å (Extended Data Fig. 14a). Although the quality of the cryo-EM map in this case was lower than for the solubilized pores, the cryo-EM density was present in the regions corresponding to the positions of phospholipids L1-5 and all four CH molecules (Extended Data Fig. 14b), proving that the cholesterol binding positions found in the Fav pores are not an artefact of pore solubilization.

In summary, SM binds specifically to positions of lipids L1-L5 in the Fav pore. Moreover, the presence of cholesterol in the membranes enables compact binding of lipids to the pore and stabilization of the pore within the membrane. This is reflected in the lower increase in ionic current noise during pore formation in cholesterol-containing membranes compared to cholesterol-free membranes (Fig. 2f and g) and in the formation of stable nonameric pores that does not dissociate during extraction from vesicles, in contrast to those formed on DOPC:SM membranes (Extended Data Fig. 7).

## Loops 2 and 3 are immersed in the headgroup region of the lipid bilayer

Once the pore is formed, the amphipathic N-terminal α1-helix of each protomer spans the membrane at an angle of ∼20° (Extended Data Fig. 15a). During pore formation, the α1-helix is extended, from I91-V100 in the soluble monomer to A80-N103 in a transmembrane protomer of the Fav pore. The α1-helix crosses the membrane with the hydrophobic side facing the membrane along its entire length. Loops 2 and 3 are immersed in the layer of phospholipid headgroups of L1, L2, L3, L4, L6 and L8, and are lined at the bottom with the cholesterol nanodomain (Extended Data Fig. 15b-d). While the overall structure of loop 3 remains virtually unchanged during the transition from the monomer in solution to the membrane-bound protomer, loop 2, residues T153-A158, moves approximately 2 Å away from the N-terminus to allow the detachment of the α1-helix from the central β-sandwich and its extension and transition through the lipid bilayer (Extended Data Fig. 16). Since the position of the two loops is independent of the presence of cholesterol, they can be described as rigid objects immersed in the headgroup region of the lipid bilayer (Extended Data Fig. 15d).

## Sphingomyelin is essential for the oligomerization of Fav

It is known that actinoporins require SM^30,31^ to effectively bind to the membrane and form pores. Our pore structures showed that lipids L1 and L6 play an important structural role. The actinoporins sticholysin II, equinatoxin II and bryoporin can bind to ceramide phosphoethanolamine (CPE), which has a smaller ethanolamine headgroup, but their permeabilization activity in CPE-containing membranes is much lower compared to SM-containing membranes^32,33^. Interestingly, we could not to obtain pores from vesicles composed of DOPC:CPE:cholesterol 1:1:1 (mol:mol:mol) (Extended Data Fig. 17a, b), using our established method for preparation of soluble Fav pores (Extended Data Fig. 1e). Cryo-EM micrographs of multilamellar DOPC:CPE:cholesterol 1:1:1 vesicles incubated with monomeric Fav showed that the membranes were completely covered with the protein, but no pores were observed. 2D class averaging showed densely packed protein particles corresponding in size to a monomeric form of Fav bound to the membranes in multiple rows (Extended Data Fig. 17b). CPE thus enables the binding of protein to the membranes, but is not capable of triggering oligomerisation.

Due to its position in the pore structure, L1 is probably the key lipid responsible for oligomerisation. The L1 head group is held in its position by formation of hydrogen bonds between its phosphate group and the side chain of R151 and the amide nitrogen of N154 of one protomer and the amide nitrogen of G243 of the adjacent protomer, and by cation-π interactions between the phosphocholine^28,34^ headgroup L1 and the Y186 side chain of the neighbouring protomer (Extended Data Fig. 10 and 18). Cation-π interactions may be significantly weaker when the phosphocholine headgroup from SM is replaced by the smaller phosphoethanolamine headgroup in CPE (Extended Data Fig. 18b). When CPE is placed at the position of L1, it additionally loses interactions with residue S241 (Extended Data Fig. 18b), which in the case of SM, altogether, stabilise the protomers in an oligomerised form (Extended Data Fig. 18).

## The Fav pore influences the bulk membrane lipids

All-atom molecular dynamics (MD) simulations of the protein pore in a POPC:SM:cholesterol 1:1:1 (mol:mol:mol) membrane were used to verify bound lipid molecules and clarify the effect of the protein pore for the surrounding lipid molecules. During the 1.5 µs MD simulation, starting from the single pore structure with 80 SM and 32 cholesterol molecules in their predefined binding positions, not a single one of these predefined lipid molecules was detected to leave its original position (Fig. 3a and b, Extended Data Fig. 19).

**Fig. 3.**
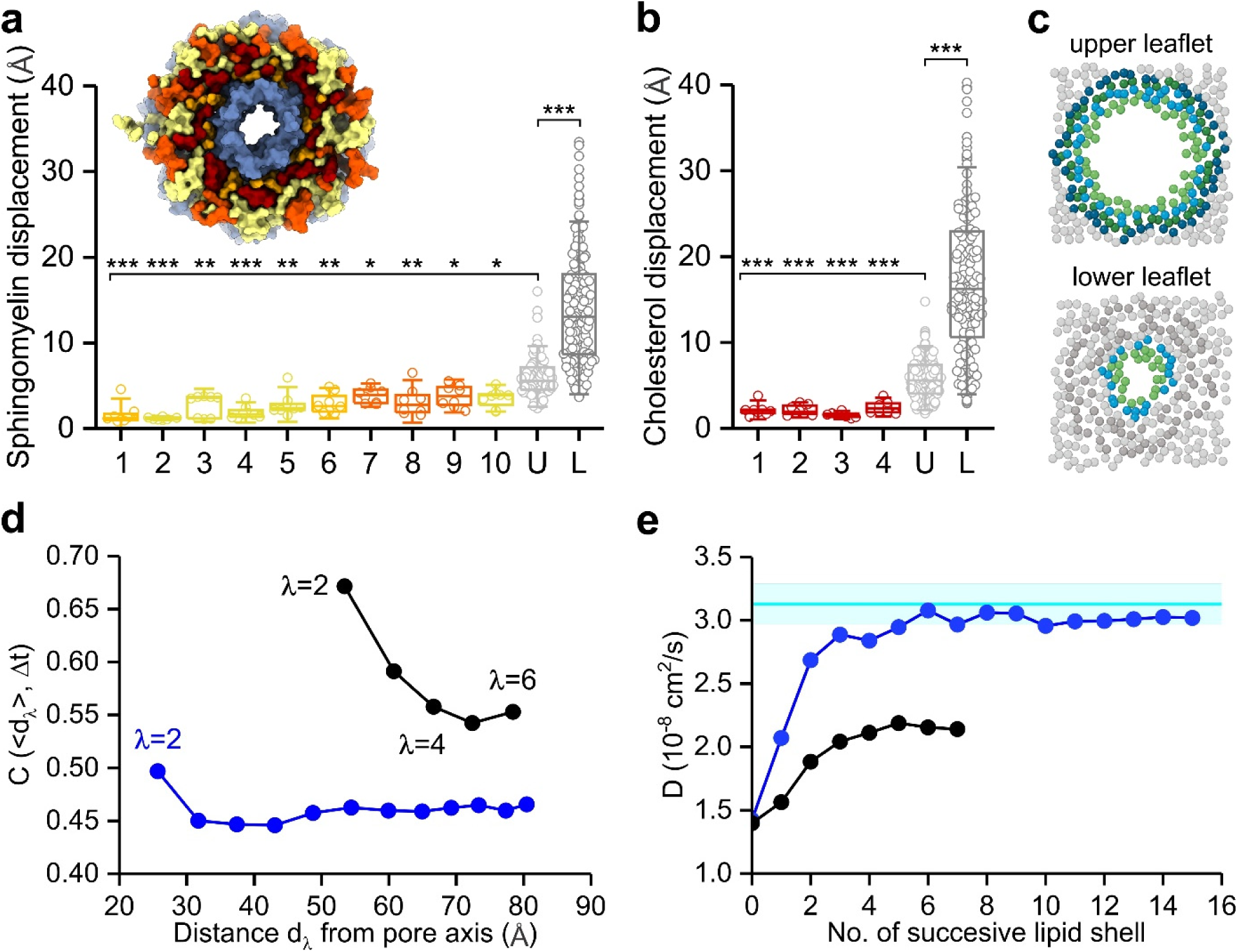
Structural and dynamic properties of membrane lipids from molecular dynamics simulations. **a**,**b** Displacement of all predetermined sphingomyelin (1-10) (**a**) and cholesterol (1-4) (**b**) lipids as well as bulk lipids in the upper (U) and lower (L) leaflet during the 1.5 μs molecular dynamics simulation. Two-sample t-test was used for statistical analysis of displacement of lipids in the upper leaflet with all other groups of lipids. * p<0.01, ** p <0.001, *** p<0.0001. The inset in **a** shows the bottom view of the pore after 1.5 μs of simulation with all associated lipids labelled according to colouring scheme in Fig. 2d. **c**, Schematic bead representation of SM molecules, which are divided into successive coordination shells in upper and lower membrane leaflet. Green colour represents lipids L3, L5, L7, L8, L9 and L10 in the upper leaflet. Each next successive shell is coloured blue and green, bulk lipids are grey. **d**, Correlation between displacements of the neighbouring lipid molecules belonging to successive shells: current (λ) versus previous shell (λ-1) of lipids for the upper (black circles) and lower leaflet (blue circles) according to equation (1). **e**, Lateral diffusion constants of the lipid molecules in successive lipid shells around the pore in the upper (black) or lower (blue) leaflet. The diffusion constants obtained for the pure membrane systems are given with the cyan line, together with the error range. Here, lipid shell 0 refers to diffusion constant due to the nearest protein monomer.

To elucidate the effect of the pore-lipid interactions on structural and dynamic features of the pore surrounding lipid molecules, we focused on the correlation between displacements of lipid molecules as well as the lateral lipid diffusion as a function of the distance from the pore. We grouped SM and POPC lipid molecules into successive coordination shells around the pore in the upper and lower leaflet of the membrane as indicated in Fig. 3c. According to the evaluated correlation, the relative motion of lipid molecules is significantly more correlated in the upper leaflet as compared to the lower one (Fig. 3d). The first lipid shell (λ=1) in both leaflets corresponds to lipid molecules in direct contact with the pore, specifically in the upper leaflet these are limited to phospholipid molecules of experimentally determined lipids involving L3, L5, L7, L8, L9 and L10. The subsequent shells (λ=2, 3…) already refer to membrane lipids interacting exclusively with lipids. As a consequence of the higher correlation driven by the strong lipid-protein cap interaction, the diffusion of lipids in the upper leaflet is significantly slower compared to the lower leaflet as shown in Fig. 3e.

## Conclusions

The cryo-EM structures of the Fav pores reveal general principles of protein interaction with lipids, such as details of interactions of protein with headgroups of sphingolipids and how lipids affect oligomerization, interaction with cholesterol and its effect on pore stability, how protein loops are immersed in the lipid bilayer, and together with MD simulations also highlight the concept of annular lipids that extend further than the first interactive shell of lipids.

Based on the structural and functional data, we can assign different functional roles to the lipids. The ***structural lipids*** L1 and L6 are integral part of the oligomeric pore structure. They provide an important interaction surface between the protomers and are thus an essential structural component of the pore. L1 is sphingomyelin, which is essential for the oligomerization of the protein protomers into a stable octameric membrane complex. One Fav protomer binds four ***receptor lipids***, L2-L5, which were also confirmed at the high-resolution level to be SM molecules, for which the protein surface is very well suited. They are stabilised by cation−π interactions at the headgroup region of the membrane and this may represent a mechanism for specific lipid recognition by membrane proteins^34^. The structures also show a cholesterol nanodomains that fit very well into the gap under the cap of the Fav pore, next to the transmembrane α1-helix, and consequently stabilize the inner and outer shell of phospholipids surrounding the pore in the upper membrane leaflet. We provide high-resolution evidence that the Fav surface can uniquely bind multiple sphingomyelin headgroups, in contrast to a single lipid receptor for peripheral membrane proteins^35,36^. The cap of the Fav pore is immersed in the lipid membrane as a rigid object via the insertion of loops 2 and 3. This means that the surface of the soluble Fav monomer is already highly adapted for membrane binding, involving multiple SM headgroups, and insertion. The advantage of binding of multiple lipid molecules is that the protein is more stably anchored to the lipid membrane. The ***bridging lipids*** have very little or no contact with the protein and can provide additional stability for pore assembly through extensive lipid-lipid interactions.

Most of the bound lipid molecules found in the cryo-EM pore structure exhibit low mobility in MD simulations, suggesting that the cap region of the pore almost completely restricts lipid movement below the cap and that the pore sits on a lipid membrane patch consisting of at least 112 lipid molecules all of which stayed at their predefined position during 1.5 µs of the simulation. The pore structure also makes it possible to study the diffusion of lipids in the two lipid membrane leaflets. In the upper leaflet, the phospholipid molecules interact with the bridging lipids, while in the lower membrane leaflet the interactions take place with the cluster of transmembrane helices. The movement of lipids estimated from atomistic simulations is correlated with the movement of the pore based on the distance to it (Fig. 3d). Interestingly, the correlation of the movement between lipids falls off differently in the two leaflets. In the bottom leaflet where we observed no specific lipid binding sites on the α-helices, the lipid-lipid correlation falls off completely after lipid interaction shell 2, whereas the lipid-lipid correlation remains significant for shells 3 and even 4 in the upper leaflet. This excludes the predefined lipids observed with cryo-EM and indicates that specific protein-lipid interactions have an ordering effect that reach further than just one additional lipid shell.

In summary, the structures of an actinoporin pore presented in this work illustrate the intricate interplay between the transmembrane protein and lipids and provide insights into the unique structural adaptations, ordering effects and functional consequences induced by specific lipid components. These findings contribute to a more comprehensive understanding of the behaviour of membrane proteins and lipids in membranes, highlighting the important structural and functional role of membrane lipids.

## Methods

### Protein expression and purification

Genes for all three Fav variants (wtFav, ΔN53Fav and ΔN75Fav with mutations R203D and D215N) were cloned into a modified pET28a plasmid, with a His6-tag followed by TEV-cleavage site preceding the N-terminus of the target protein. Transformed *E. coli* BL21(DE3) cells were grown at 37 °C in Terrific Broth (TB), and gene expression was induced with 0.4 mM isopropyl β-D-1-thiogalactopyranoside when A_600_ reached around 0.7. After induction, the temperature was lowered to 20 °C. Cells were harvested after 16 hours by centrifugation at 6,000 × *g* for 5 min at 4°C, followed by sonication in PBS buffer pH 7.4 (1.8 mM KH_2_PO_4_, 140 mM NaCl, 10.1 mM Na_2_HPO_4_, 2.7 mM KCl). The cell lysate was centrifuged at 50,000 *g*, 4 °C for 45 min. The supernatant was filtered using a 0.22 μm syringe filter and loaded onto a NiNTA 10/50 column (Qiagen, Germany) and proteins were eluted with an increasing gradient of imidazole. The haemolytic fractions were pooled and incubated over night with addition of TEV protease with a final concentration of 1% (w/w) at 20 °C. After imidazole was removed with dialysis against PBS, the protein was again loaded onto a NiNTA 10/50 column (Qiagen, Germany) and subsequently subjected to size exclusion chromatography using Superdex 200 prep grade column (GE Healthcare, UK) equilibrated with PBS. Protein-containing factions were concentrated with Amicon Ultra Filter Devices (Millipore, USA) 10 kDa cutoff, aliquoted and stored at - 70 °C.

### Protein crystallization and crystal structure determination

High-quality ΔN53Fav monomer crystals were obtained by mixing 1 µl of the protein solution with a concentration of 30 mg/ml with 1 µl of the reservoir solution containing 1.8 M Li_2_SO_4_ using the vapor-diffusion technique in hanging drops. The drop was equilibrated at 20 °C over 0.5 ml of reservoir solution. The rod-like crystals typically appeared within two days. Crystals were frozen in liquid nitrogen, with 20 (v/v) % 2-methyl-2,4-pentanediol as a cryoprotectant. Diffraction data was collected at 100 K and at the wavelength of 1.0 Å at XDR1 Elettra Synchrotron (Trieste, Italy). The diffraction data was processed to 1.5 Å resolution with XDS^37^ (Extended Data Table 1). The crystal structure was solved using the symmetry of the space group P3_2_21 by molecular replacement (PHASER^38^), with the crystal structure of FraC (PDB-ID 3ZWJ) without loops and α-helices as a search model. Initial Δ53 Fav model was constructed with PHENIX^39^ Autobuild and refined by iterative cycles of manual model building in Coot^40^ and phenix.refine^41^. The crystal structure of Δ53 Fav is missing the first 25 N-terminal residues of Δ53Fav construct. There are regions of continuous electron density than likely correspond to the N-terminal region, but the density was too weak to be interpretable. The overall structure has great geometry with Molprobity score of 1.1 and the following Ramachandran parameters: favored (98.35%), allowed (1.65%), and outliers (0.00%). The graphical presentation of the structure in the figure was done using ChimeraX v1.7^42^.

### Preparation of lipid vesicles

Lipid vesicles were prepared using lipids from Avanti Polar Lipids, USA. CPE, DOPC, POPG, SM, and cholesterol were dissolved in chloroform or methanol and mixed at appropriate molar ratios (1:1 or 1:1:1). Using a rotavapor (Büchi, Switzerland), a thin lipid film was created and left under a high vacuum for 2 hours. The multilamellar vesicles were produced by resuspending the lipid film in PBS buffer and thoroughly vortexed with the aid of 0.5 mm glass beads (Scientific Industries, USA). The vesicles suspension then underwent at least three fast freeze/thaw cycles. Unilamellar vesicles were formed by passing the multilamellar vesicles through a LiposoFast lipid extruder (Avestin, Canada) with polycarbonate membranes with 100 nm pores.

### Pore preparation

70 µM of the wild-type Fav monomer was incubated with 14 mM large unilamellar vesicles in PBS buffer for 1 h at 37 °C. The mixture was solubilized with 1 % lauryldimethylamine oxide. Sample was centrifuged for 5 min at 16,000 *g* at 20 °C to remove precipitated material. Supernatant was 20-fold diluted in buffer A (50 mM Tris, 0.25 mM Brij 35, pH 7.4) and injected to the Resource Q column (Pharmacia Biotech, Sweden) equilibrated with buffer A. Proteins were eluted with a linear gradient of buffer B (50 mM Tris, 1 M NaCl, 0.25 mM Brij 35, pH 7.4). Oligomeric state of proteins in eluted fractions was monitored with Native PAGE.

### Nanodisc preparation

Plasmid pMSP1E3D1, bearing the gene for membrane scaffold protein expression was a gift from Stephen Sligar (Addgene plasmid # 20066) and was expressed as described before^43^. Shortly, *E. coli* BL21(DE3) cells transfected with the plasmid were grown in TB medium at 37 °C until A_600_ reached 2.5. Protein production was initiated with the addition of 1 mM isopropyl β-D-1-thiogalactopyranoside. Cells were harvested 3.5 h later, centrifuged at 6000 *g* and the pellet stored at – 80 °C. MSP1E3D1 protein was purified using Chelating Sepharose FF (GE Healthcare, UK) and dialyzed against MSP standard buffer (20 mM TRIS-HCl, 100 mM NaCl, 0.5 mM EDTA, pH 7.4). Protein was concentrated with Amicon Ultra-15 (Millipore, USA) to a final concentration 9 mg/ml.

The assembly of lipid nanodisc was carried out as described^44^ with the following adjustments. A mixture of 5 mg DOPC, 5 mg SM and 2.5 mg cholesterol was dissolved in chloroform and methanol 1:1 (volume) mixture, and then dried in a rotary evaporator Rotavapor R215 (Buchi, Switzerland). A thin lipid film, formed in a round-bottom flask, was then hydrated by the addition of 100 µl of cholate buffer (20 mM Tris, 25 mM Na-cholate, 140 mM NaCl, pH 7.4), and vigorous vortexing and heating using warm tap water. The mixture was then sonicated in a water bath for 15 min. Finally, 250 µl of purified MSP1E3D1 were added and the mixture was incubated on a bench shaker at 4°C for 3 h. The nanodiscs were self-assembled upon removal of Na-cholate by overnight dialysis against 3 l of PBS buffer. The dialyzed sample was purified using size-exclusion chromatography (Superdex 200 10/300, GE Healthcare Life Sciences, USA). Peak fractions, corresponding to the size of nanodiscs, were analyzed with dynamic light scattering with Prometheus Panta (Nanotemper, Germany) to confirm the presence of ∼12 nm particles and the homogeneity of the solution. Concentration of nanodiscs was estimated by measuring the concentration of MSP at A_280_. Nanodiscs were used immediately for cryo-TEM experiments.

### Planar lipid membranes experiments

Experiments were performed with the Orbit Mini (Nanion Technologies, Germany) set up using MECA 4 chips (Ionera, Germany). The data were collected with the Elements Data Reader v 3.8.3 (Elements, Italy) software at 20 nA working range, 20 kHz sampling frequency and room temperature. Planar lipid bilayers were prepared with the brushing method, where 10 mg/ml lipid solution in pure octane was used. To achieve pore insertion, a monomeric Fav variant (deletion of 75 amino acids from the N-terminus and R203D, D215N mutations) was applied at 1.6 µg/ml final concentration to DOPC:SM 1:1 or DOPC:SM:Cholesterol 1:1:1 membranes. For statistical analysis we calculated the mean current and its standard deviation (unfiltered current traces) for the intact membrane and the first three observed insertion steps (smaller than 200 pA). NR ratio represents the ratio between the noise of the specific insertion step (standard deviation, STD) and intact membrane noise (STD)^45^. Data were analyzed using Axon pCLAMP 11.1 (Molecular Devices, USA) software and our own MatLab R2022a (Mathworks, USA) script. Plotted traces were filtered using MatLab lowpass function with cut off frequency of 1000 Hz.

### Cryo-electron microscopy sample preparation and data acquisition

Isolated wild-type Fav pores were prepared as described above (extended Figure 1d). Samples of wild-type Fav pores on vesicles were prepared similarly, excluding the solubilization and purification steps. Instead, the sample was diluted after the incubation so that the final lipid concentration was 2 mM. Wild-type Fav pores on nanodiscs were prepared by incubating nanodiscs (0.4 mg/ml) with monomeric Fav (1 mg/mg) for 1 h at 37°C. After incubation, the sample was 20 × concentrated with an Amicon Ultra Filter Device (Millipore, USA) (100 kDa cut-off) and 20 × diluted with PBS. This step was repeated three times to remove unbound monomers from the sample.

3 µl of each sample (purified pores, pores on vesicles and pores inserted in nanodiscs) was applied to glow-discharged (GloQube® Plus, Quorum, UK) Quantifoil R1.2/1.3 or R2/2 mesh 200 copper holey carbon grid (Quantifoil, Germany), blotted under 100% humidity at 4°C for 6–8 s, and plunged into liquid ethane with a Mark IV Vitrobot (Thermo Fisher Scientific, USA). Micrographs of the wild-type Fav pore prepared on DOPC:SM vesicles were recorded on Titan Krios G2 (Thermo Fisher Scientific, USA) operated at 300kV with K2 direct electron detector (Gatan, USA) at Diamond Light Source (UK), all other samples were recorded inhouse on Glacios (Thermo Fisher Scientific, USA) with a Falcon 3EC direct electron detector (Thermo Fisher Scientific, USA) and operated at 200 kV using the EPU software (Thermo Fisher Scientific, USA).

### Cryo-electron microscopy data processing

All steps of data processing for all samples were performed in cryoSPARC v4^46^ with built-in algorithms. Movies were aligned and dose-weighted with patch motion correction, CTF was estimated with patch CTF. Micrographs with estimated CTF fit estimation above 6 Å were excluded from further analysis. Initially, particles were picked by hand to generate 2D templates for template picking. From this point on the analysis slightly diverged for the three samples but followed similar steps. Workflows for each sample are provided (Extended Data Fig. 2, 7, 13, and 14). Particles were extracted and underwent several rounds of 2D classification. Particles from best classes were used in *ab-initio* reconstruction. Particles were then further cleaned up by heterogeneous refinement. Particles from the best class were then re-extracted using local motion correction and used in homogeneous refinement and non-uniform refinement with applied C8 symmetry.

### Model building

Atomic models of protomers were built with the iterative use of Coot^40^ and Isolde^47^ plugin for ChimeraX^48^. Phenix^49^ was used for applying symmetry and restraint calculation for lipid molecules. The crystal structure of ΔN53Fav monomer was used as the starting model. Details of data acquisition and refinement statistics are shown in Extended Data Table 3. The graphical presentation of the structures and cryo-EM maps in figures was done using ChimeraX v1.7^42^.

### Molecular dynamics simulations

All-atom models of the protein pore were prepared using the online server CHARMM-GUI^50^. Lipid bilayers composed of POPC:SM:Cholesterol 1:1:1 (mol:mol:mol) were used. The cryo-EM protein pore structure containing 80 predefined SM and 32 cholesterol molecules was inserted into the POPC:SM:Cholesterol membrane. Membrane composed of POPC:SM:Cholesterol 1:1:1 (mol:mol:mol) was used to assess the diffusion of the lipids in the absence of the Fav pore. Both systems were electro-neutralized and immersed in 150 mM NaCl aqueous solution within the simulation box of dimensions 145 Å × 145 Å × 138 Å.

Each system was exposed to minimization and to a long equilibration phase of 200 ns. All simulations were carried out on GPU’s with the CUDA version of the NAMD molecular dynamics software suite^51^. The CHARMM36 force field^52^ was used and water was modelled by the TIP3P water model^53^. All production simulations used the NPT ensemble and their length was at least 1 µs. Temperature was held constant at 303.15 K using the Langevin thermostat with a dampening constant of 1 ps-1. The pressure was held constant at 1.0 bar. The cut-off for nonbonded interactions was set to 12 A, electrostatic interactions were calculated using the Particle Mesh Ewald method (PME).

The individual lipid displacements are calculated as the average distance travelled by each lipid within 1 microsecond. Correlation between displacements of molecules I and J, which are represented by centres of mass of selected atoms (beads), was evaluated through the ensemble average of displacements taking place during the time *Δt*:

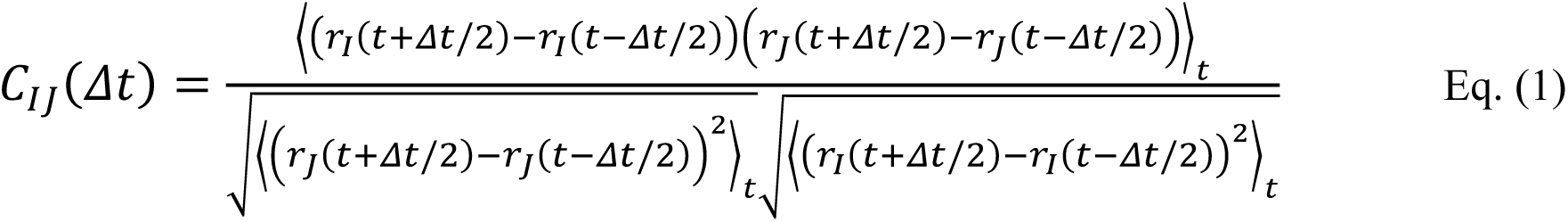

Displacements during the time step *Δt*=5 ns were used in correlation evaluation. Further dynamic properties of the lipid molecules were analysed by evaluating lipid lateral diffusion using the Einstein relation^54^ by estimating the slope of the linear fit to the time dependence of the lipid molecules’ mean square displacements.

## Data availability

All models and density maps have been deposited in the PDB and the EMDB. The PDB IDs and associated EMDB ISs are 9EYL (monomeric ΔN53Fav), 9EYM and EMD-50057 (Fav pore extracted from DOPC:SM membrane), 9EYN and EMD-50060 (Fav pore extracted from DOPC:SM:cholesterol membrane), 9EYO and EMD-50059 Fav pore extracted from POPG:SM:cholesterol membrane), EMD-50060 (Fav pore prepared on DOPC:SM:cholesterol nanodiscs).

## Acknowledgements

This study was supported by the Slovenian Research Agency (Program grant P1-0391and projects J4-8225 and J4-2547) and Oxford Nanopore Technologies, UK. Structural characterization of Fav pores was performed at the National Institute of Chemistry Cryo-EM Facility, supported by ARIS infrastructure program IO-0003. MD simulations and part of cryo-EM data analysis was performed at the Ažman Computing Center (HPC ARC) at the National Institute of Chemistry. We also thank the Centre of Excellence for Integrated Approaches in Chemistry and Biology of Proteins (CIPKeBiP), Ljubljana, for the initial crystallization screening for ΔN53Fav monomer as well as the staff at the Elettra synchrotron, Trieste, Italy, for the valuable technical help with X-ray diffraction data collection. The authors gratefully acknowledge the HPC RIVR consortium (www.hpc-rivr.si) and EuroHPC JU (eurohpc-ju.europa.eu) for funding this research by providing computing resources of the HPC system Vega at the Institute of Information Science (www.izum.si). Part of cryo-EM data analysis was performed at the HPC cluster Arnes (www.arnes.si). We thank Tomaž Švigelj for the help with protein isolations and pore preparations. We are grateful to Matic Kisovec, Toshihide Kobayashi and Alessandra Magistrato for critical reading of the manuscript.

## Author information Authors and Affiliations

Department of Molecular Biology and Nanobiotechnology, National Institute of Chemistry, Ljubljana, Slovenia

Gašper Šolinc, Marija Srnko, Ana Crnković, Mirijam Kozorog, Marjetka Podobnik, Gregor Anderluh

Theory Department, National Institute of Chemistry, Ljubljana, Slovenia

Franci Merzel

## Contributions

Experiments were designed and conceived by G. Š., M. S., F. M., A. C., M. K., M. P., and G. A. Fav protein expression and pore preparation experiments were performed by G. Š., M. S. and A. C. Protein crystallography was performed and analysed by G. Š. and M. P. Cryo-EM structural experiments and analyses were performed by G. Š. and M. P. MSP for nanodiscs was prepared by M. K. M. S. performed planar lipid bilayer experiments and data analysis. F. M. performed MD simulations and analysed the data. M. P. and G. A. supervised the work. G. Š., M. P. and G. A. wrote the manuscript. All of the authors contributed to editing the manuscript, and support its conclusions.

## Extended Data Figure 1

**Extended Data Fig. 1.**
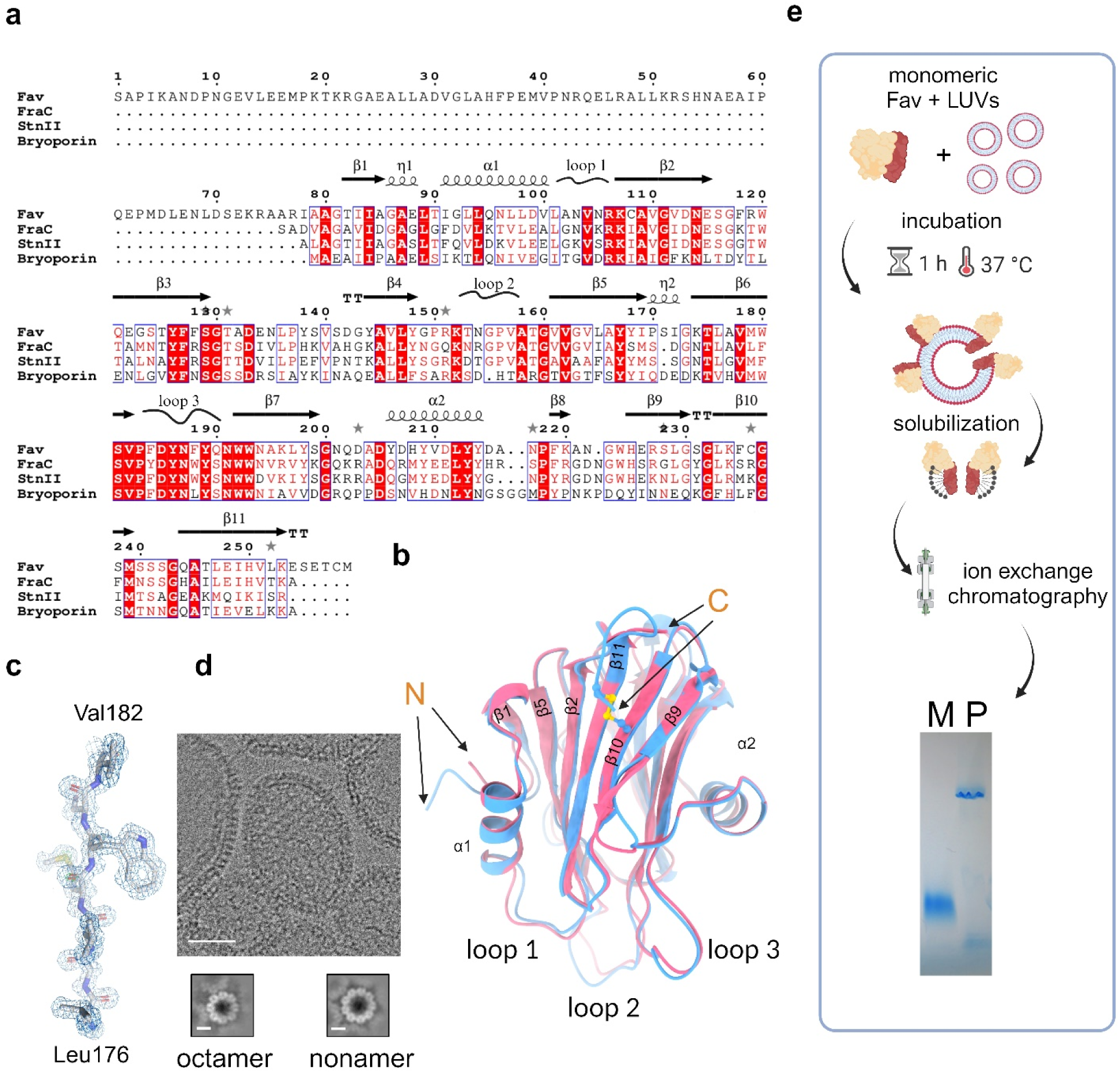
Structure of monomeric ΔN53Fav and preparation of isolated soluble wild type Fav pores. **a**, Amino acid sequence alignment of Fav with actinoporins (Fragaceatoxin C (FraC), Uniprot: B9W5G6; Sticholysin II (StnII), Uniprot: P07845; Bryoporin, Uniprot: Q5UCA8). Identical and chemically similar residues are marked with red background and red letters, respectively. The scheme of the secondary structure of ΔN53Fav is indicated above the alignment. The η symbol refers to a 3_10_-helix. α-helices and 3_10_-helices are displayed as helical lines, loops as squiggles, β-strands are rendered as arrows, and strict β-turns as TT letters. Residues with alternative conformations are indicated with grey stars above. **b**, Superposition of crystal structures of monomeric ΔN53Fav (this work, PDB-ID 9EYL, blue) and FraC (PDB-ID 3LIM, pink). N- and C-termini and secondary structure elements are labelled. The disulfide bond in ΔN53Fav between C236 and C258 is shown as yellow sticks. **c**, A structural detail in ΔN53Fav crystal structure, region L179-V185. 2mFoDFc electron density is contoured at 1 σ (blue) and mFo-DFc electron density at +3.0 σ (green) and at –3.0 σ (red). **d,** Cryo-EM micrograph of DOPC:SM 1:1 (mol:mol) large unilamellar vesicles (LUVs) incubated with monomeric wild type Fav (top) and 2D class averages of two different pore stoichiometries. Scale bars represent 50 preparation steps, from top to bottom: monomeric Fav was mixed with LUVs (blue circles) (not drawn to scale), and incubated for1 h at 37°C. Membranes were then solubilized with 1% lauryl dimethylamine oxide. Extracted Fav pores were further purified using ion exchange chromatography and analyzed by Native-PAGE (left: monomeric Fav (M), right: purified Fav pore (P)).

**Extended Data Fig. 2.**
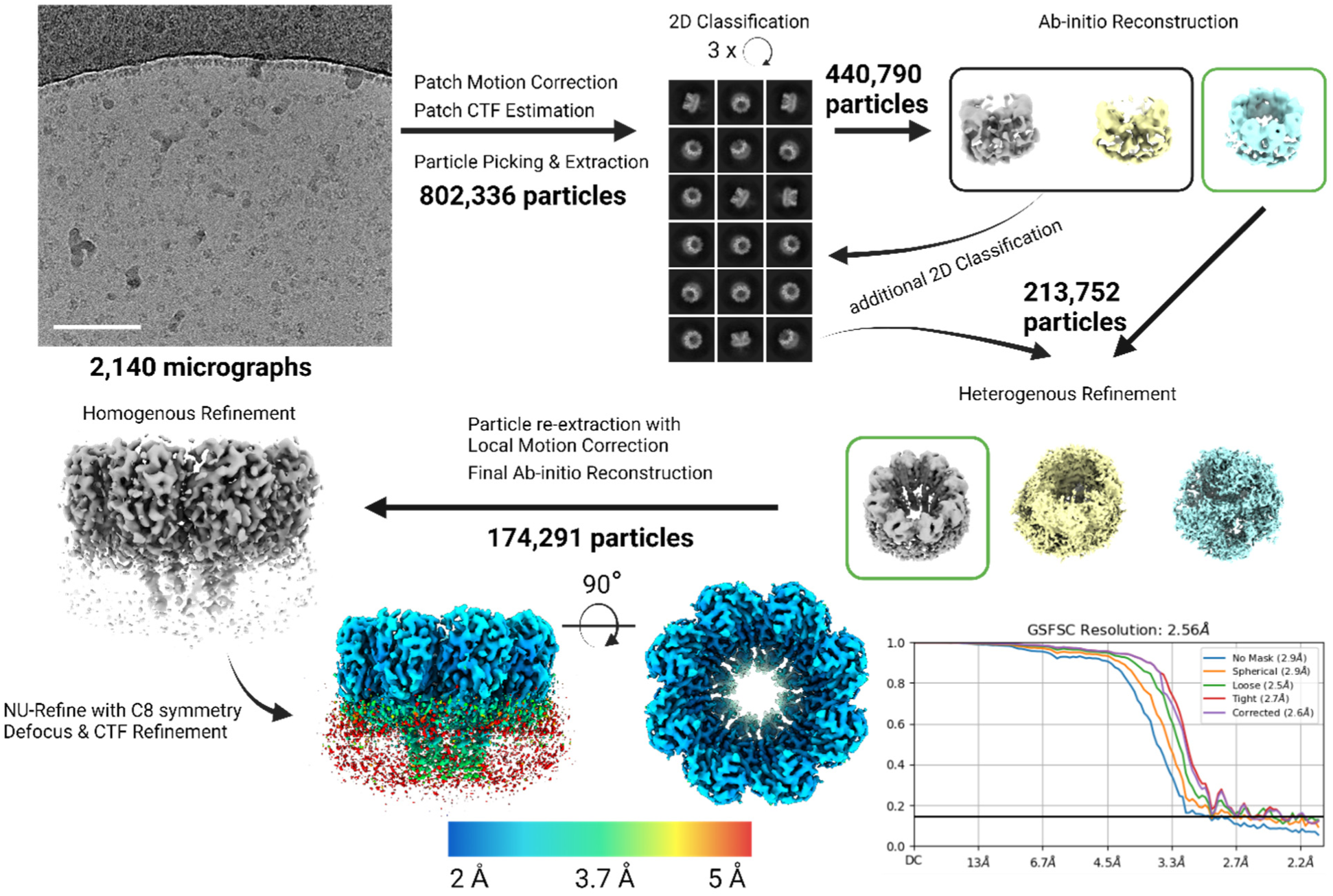
Cryo-EM data analysis workflow for isolated wild type Fav pores extracted from large unilamellar vesicles composed of DOPC:SM 1:1 (mol:mol). Additional details on data analysis with CryoSPARC v4^46^ are provided in cryo-EM data processing section of Materials and methods.

**Extended Data Fig. 3.**
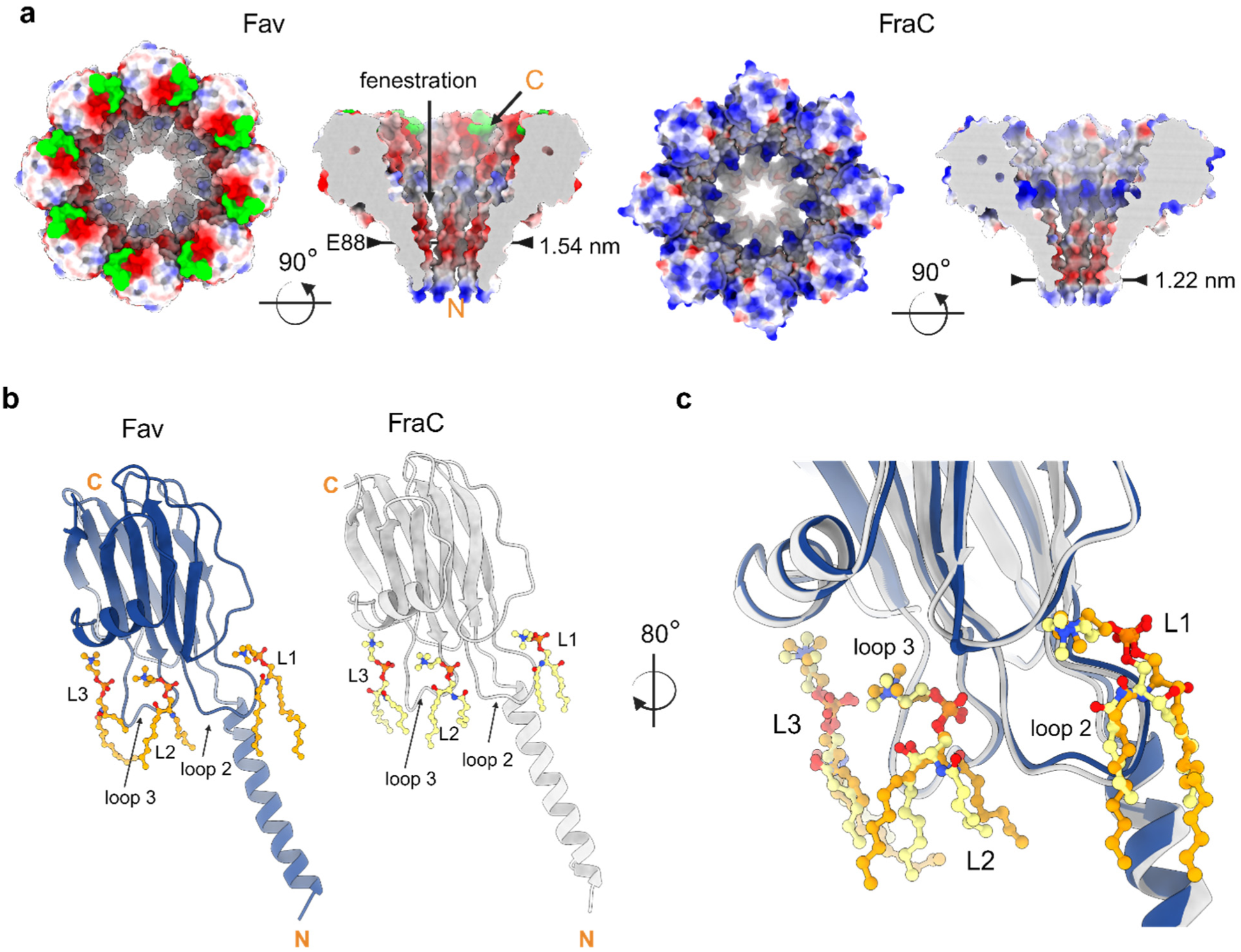
Comparison of Fav pore extracted from DOPC:SM 1:1 (mol:mol) membranes (PDB-ID 9EYM) with the crystal structure of FraC pore (PDB-ID 4TSY). **a**, The electrostatic surface potential (+/-10 kT/e) of the Fav pore and the FraC pore. The top view and cross-section are shown side by side, with the color gradient from negative (red) to positive (blue) potential. The pore constriction is highlighted with arrows and the diameter is indicated. The C-terminal extension of Fav is highlighted in green and fenestrations are indicated by an arrow. **b, c,** The comparison of lipids previously described in the crystal structure FraC pore and lipids at equivalent positions in the structure of the Fav pore. **b**, Side-by-side comparison of individual protomers with lipids L1, L2, and L3. **c**, Fav protomer (blue) aligned to FraC protomer (grey) with corresponding lipids in ball-and-stick presentation (orange carbon atoms for Fav and yellow for FraC). The C and N-terminals are marked by orange letters C and N.

**Extended Data Fig. 4.**
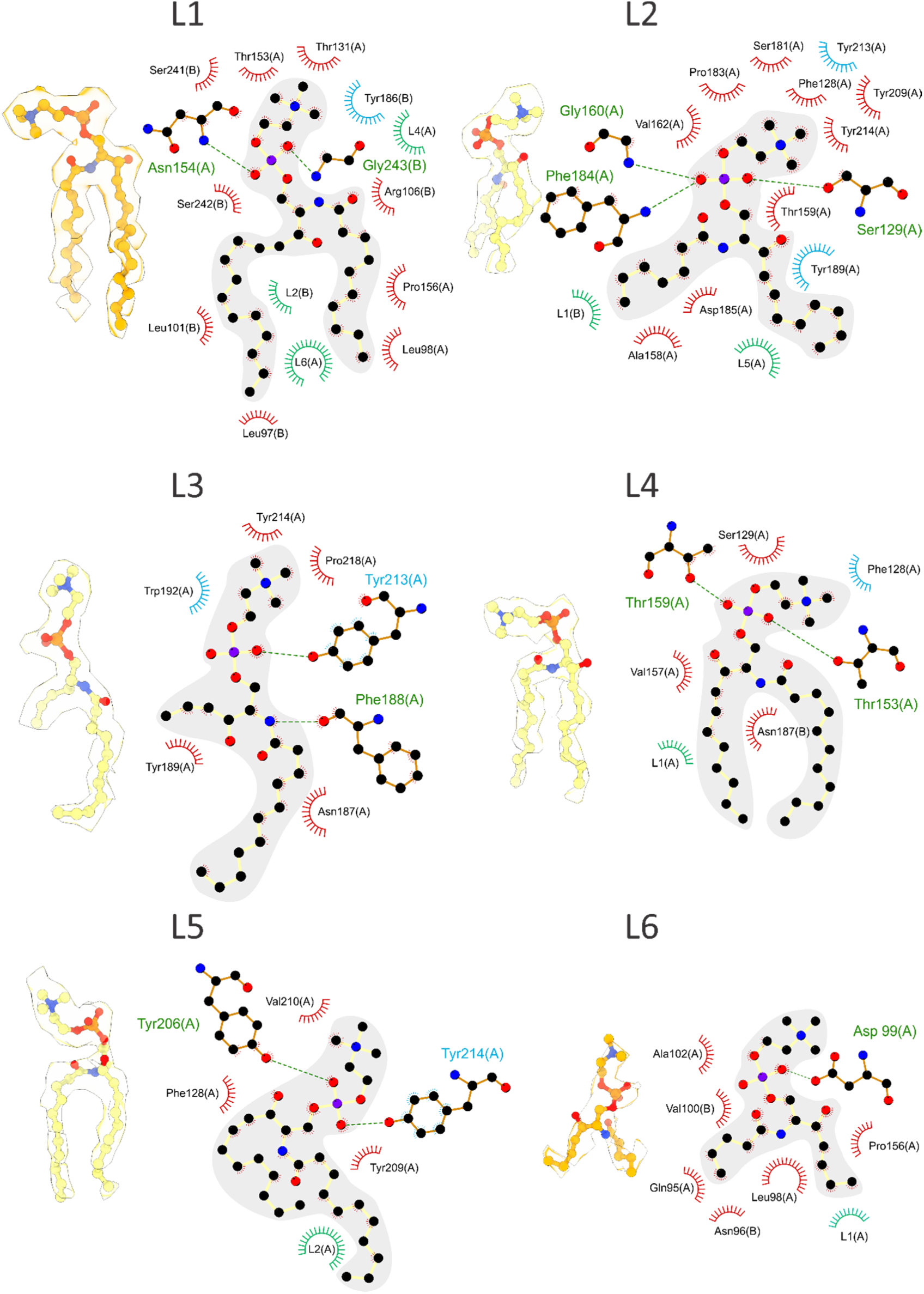
Lipids in the Fav pore structure extracted from large unilamellar vesicles composed of DOPC:SM 1:1 (mol:mol). Cryo-EM density maps with fitted models of lipid molecules are shown on the left side for each lipid, with an interaction 2D diagram on the right side. Hydrogen bonds are represented with green dashed lines. Spoked arcs and circles represent residues (red) and lipids (green) making nonbonded contacts with the lipid. Residues with name in blue or surrounded by blue arcs show cation-π interactions with the choline group. The interaction diagrams were prepared by LigPlot+^55^ and modified by hand after manual inspection.

**Extended Data Fig. 5.**
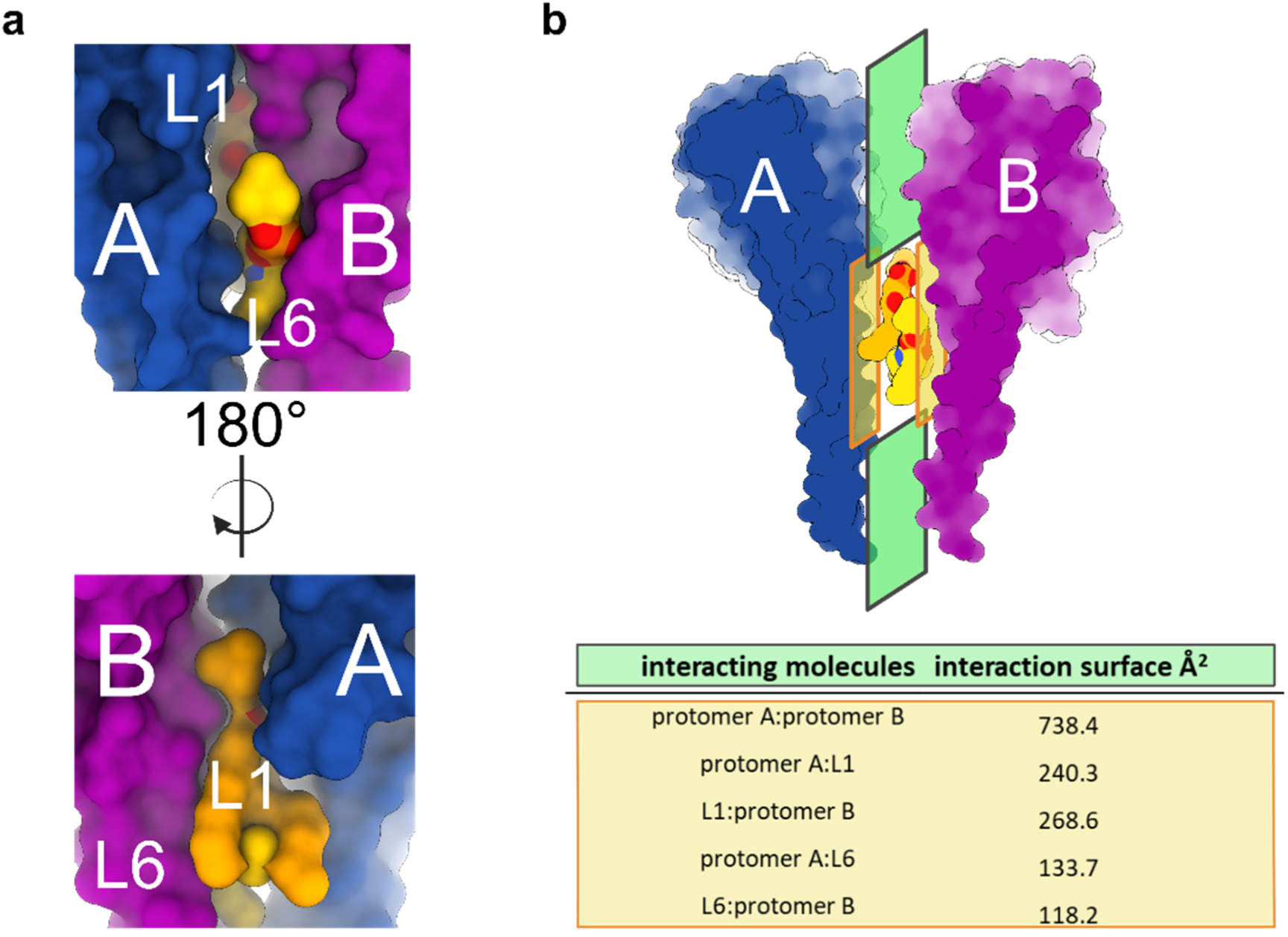
Interaction surface between two protomers in a pore. **a**, A position of lipids L1 and L6 between the protomers A and B, all molecules presented by the molecular surfaces. **b**, A schematic representation of interaction surfaces between protomers A and B. Green rectangle represents the direct protomer-protomer surface and orange rectangle the indirect protomer-lipid-protomer interactions. The table indicate interaction surfaces as calculated by pdbePISA^56^.

**Extended Data Fig. 6.**
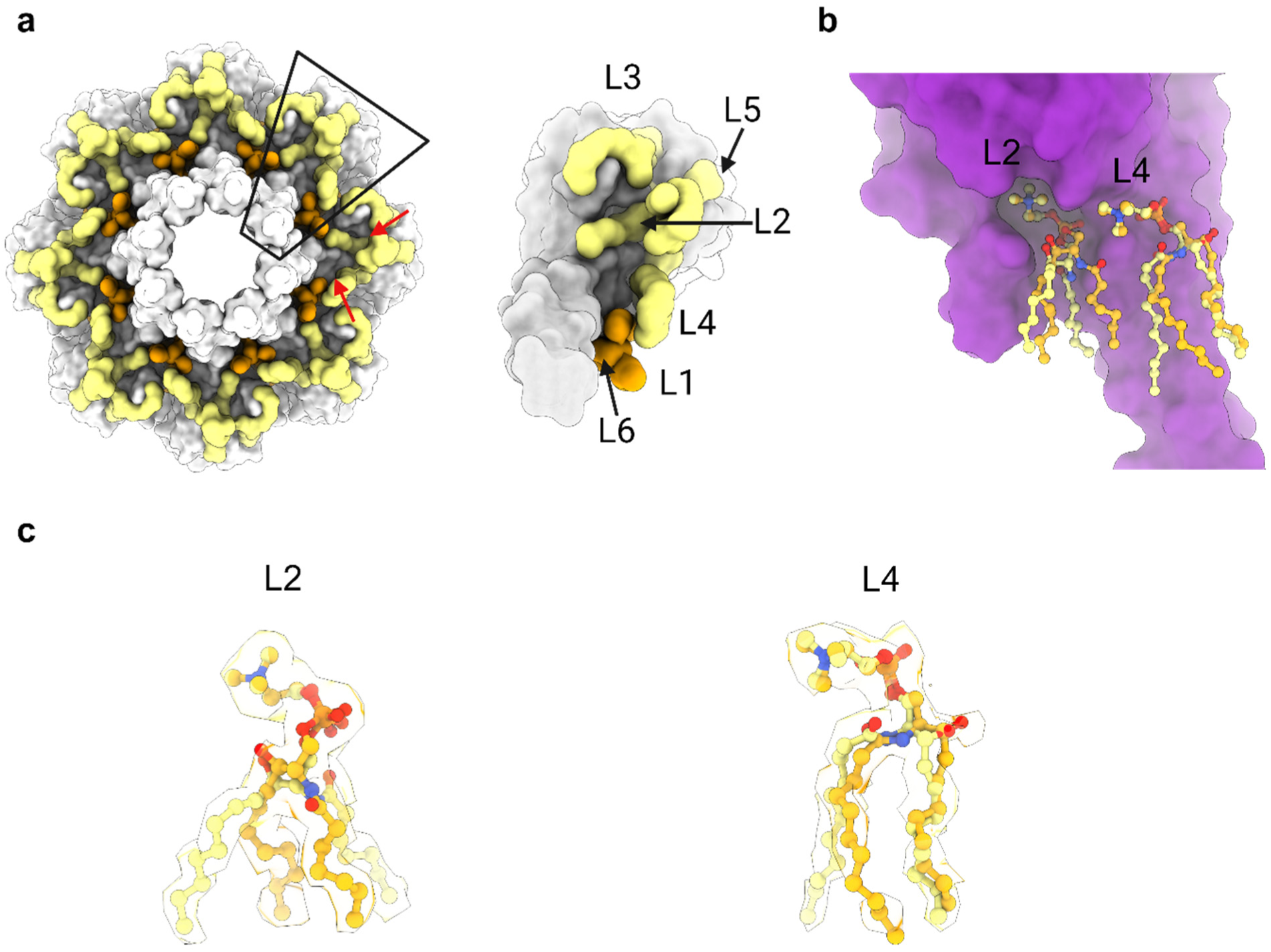
Alternative lipid chain conformations of lipids L2 and L4. **a**, bottom-up view at the pore (white) and lipids in orange and wheat. A single protomer with associated lipids is highlighted (trapezoid, black line). Red arrows show the empty space underneath the pore cap. **b**, Alternative conformations of lipids L2 and L4. **c**, Cryo-EM maps corresponding to lipids L2 and L4, filled with ball-and-stick models of lipids, highlighting alternative conformations of lipids.

**Extended Data Fig. 7.**
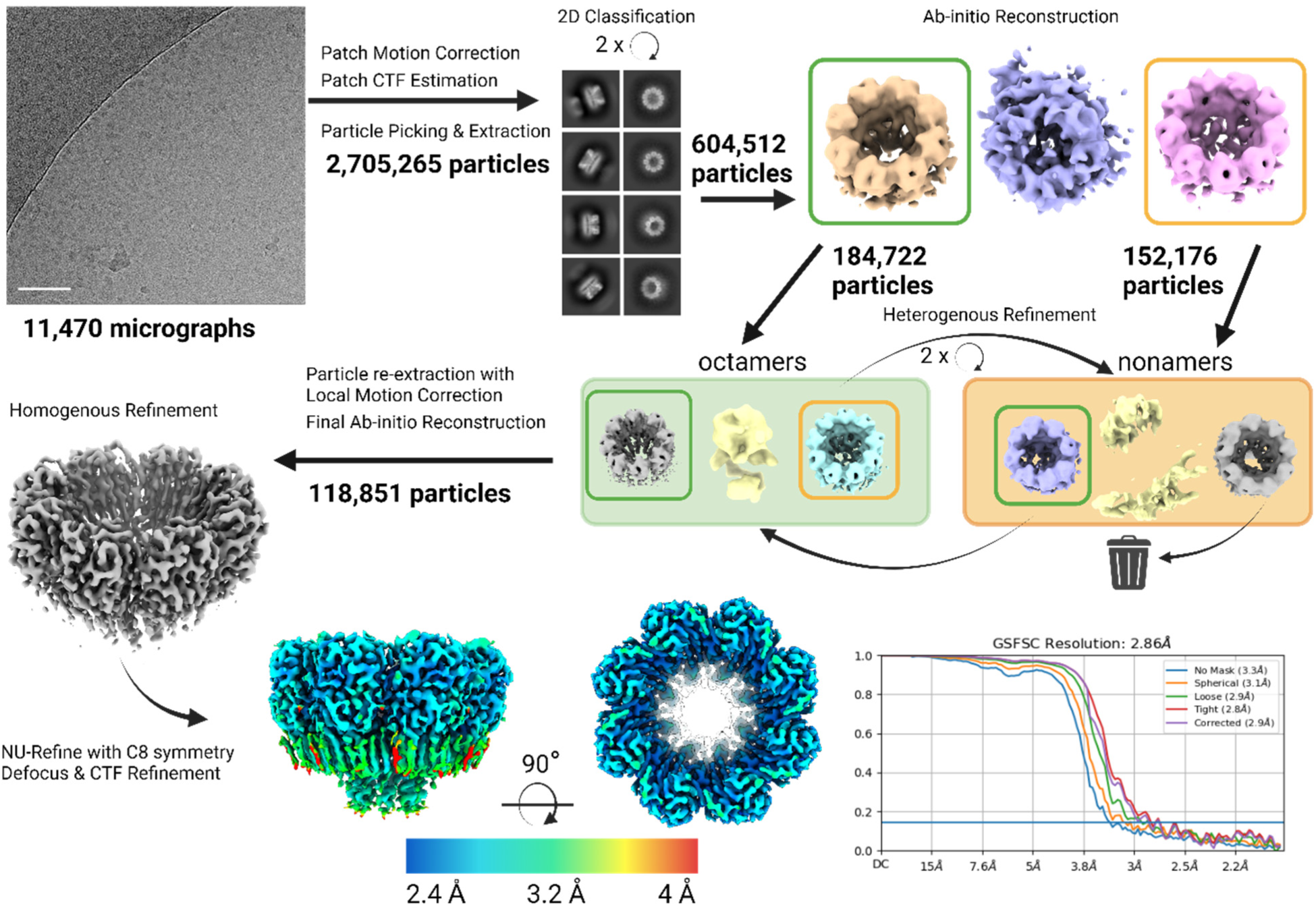
Data analysis workflow for wild type Fav pore extracted from large unilamellar vesicles composed of POPG:SM:cholesterol 1:1:1 (mol:mol:mol). All steps were performed in CryoSPARC v4^46^. Octameric and nonameric particles were separated with heterogeneous refinement after witch the nonameric particles were excluded from further analysis. Scale bar is 50 nm. Additional details on data analysis are provided in cryo-EM data processing section of Materials and methods.

**Extended Data Fig. 8.**
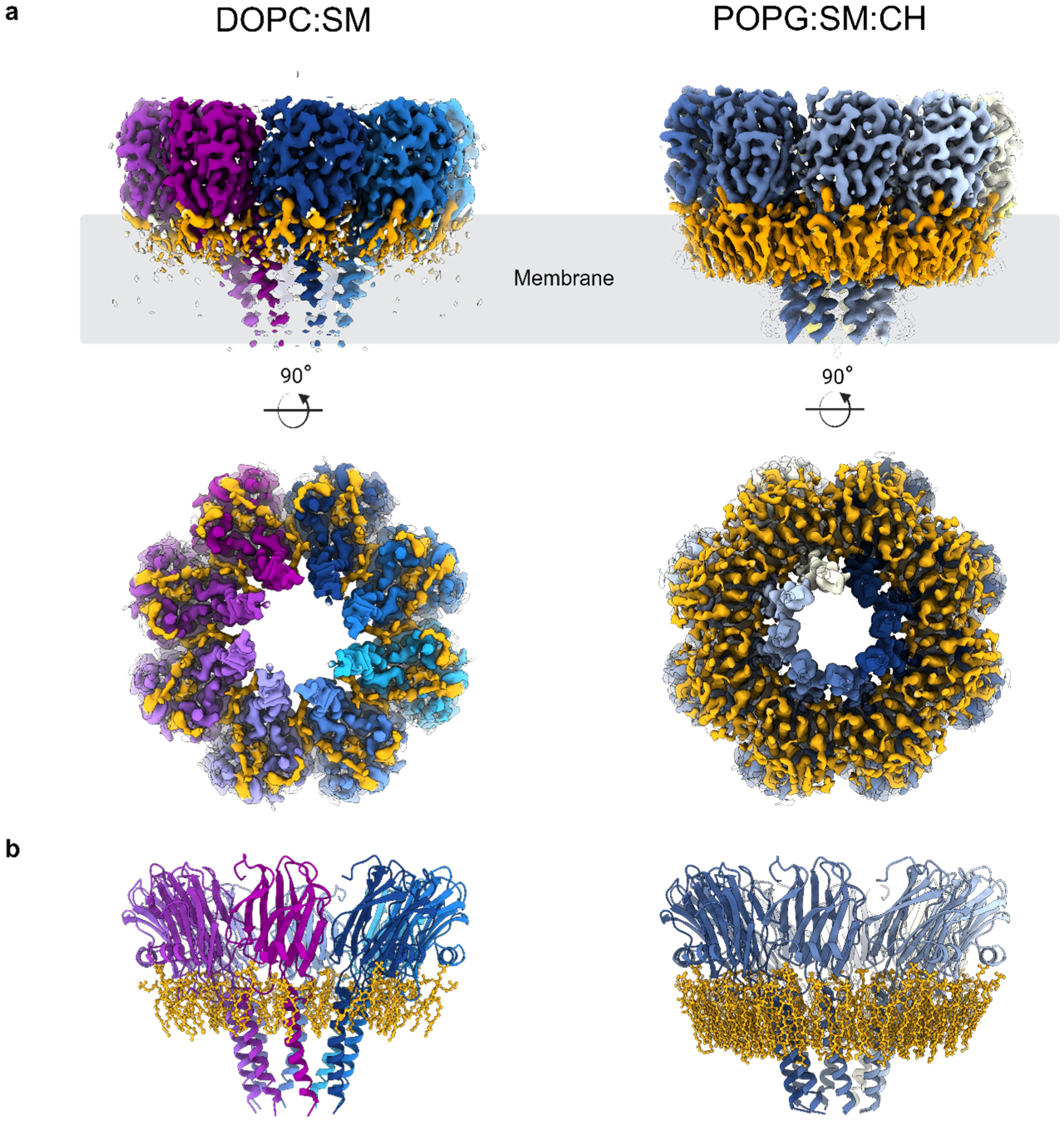
Comparison of pores extracted from large unilamellar vesicles composed of DOPC:SM 1:1 (mol:mol) or POPG:SM:cholesterol 1:1:1 (mol:mol:mol). **a**, Side and bottom views of the pore densities. Cryo-EM density maps are colored by protomers (blue and purple shades), and regions corresponding to bound lipids are shown in orange. The grey rectangle indicates the position of the lipid membrane. **b**, Models of pores with associated lipids.

**Extended Data Fig. 9.**
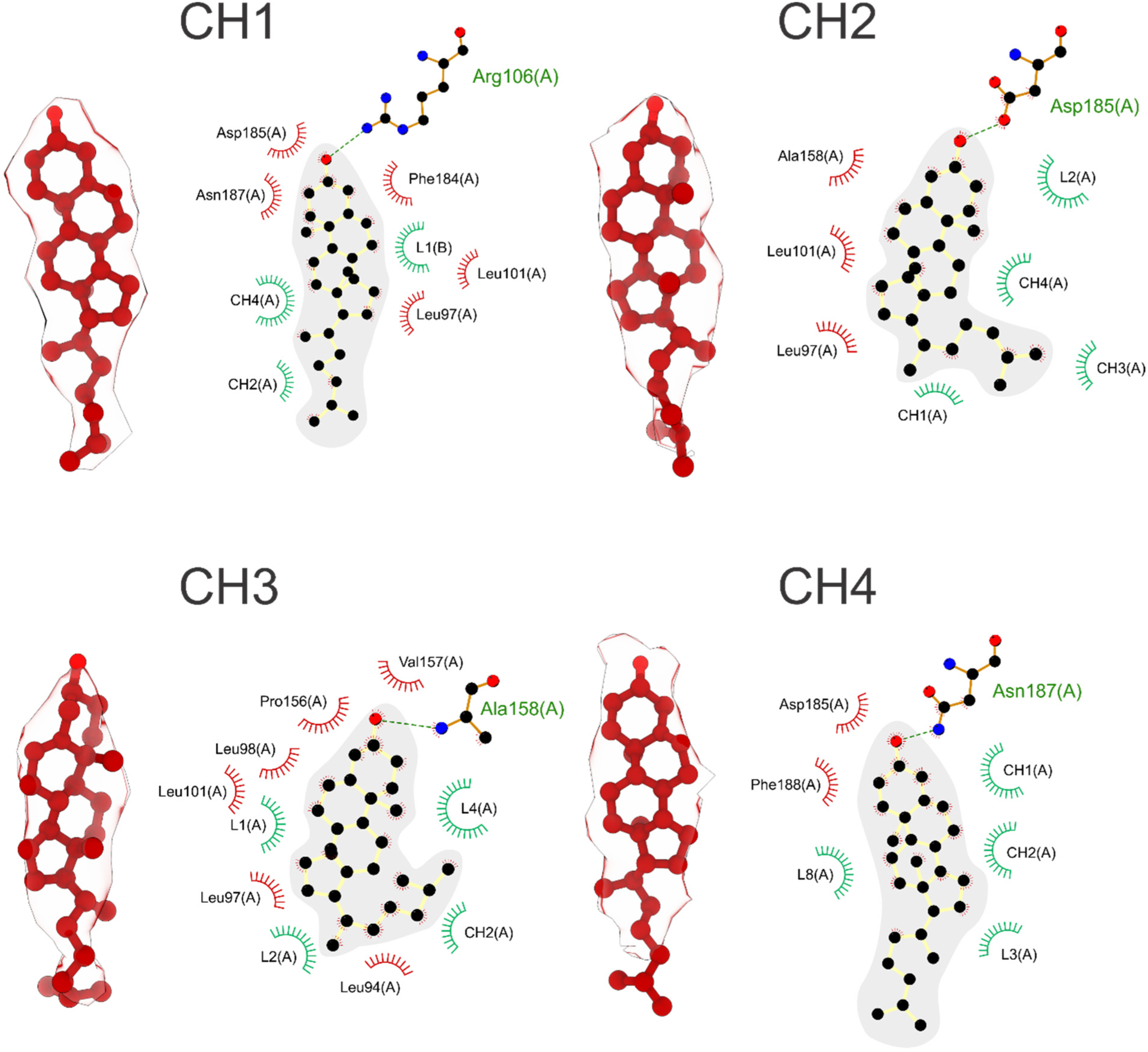
Cholesterol molecules in the Fav pore structure formed on large unilamellar vesicles composed of POPG:SM:cholesterol 1:1:1 (mol:mol:mol). Cryo-EM density maps with fitted ball-and-stick models are shown for each cholesterol molecule (left), with an interaction 2D diagram (right). Hydrogen bonds are represented with green dashed lines. Spoked arcs and circles represent residues (red) and lipids (green) making nonbonded contacts with the lipid. The interaction diagrams were prepared by LigPlot+^55^.

**Extended Data Fig. 10.**
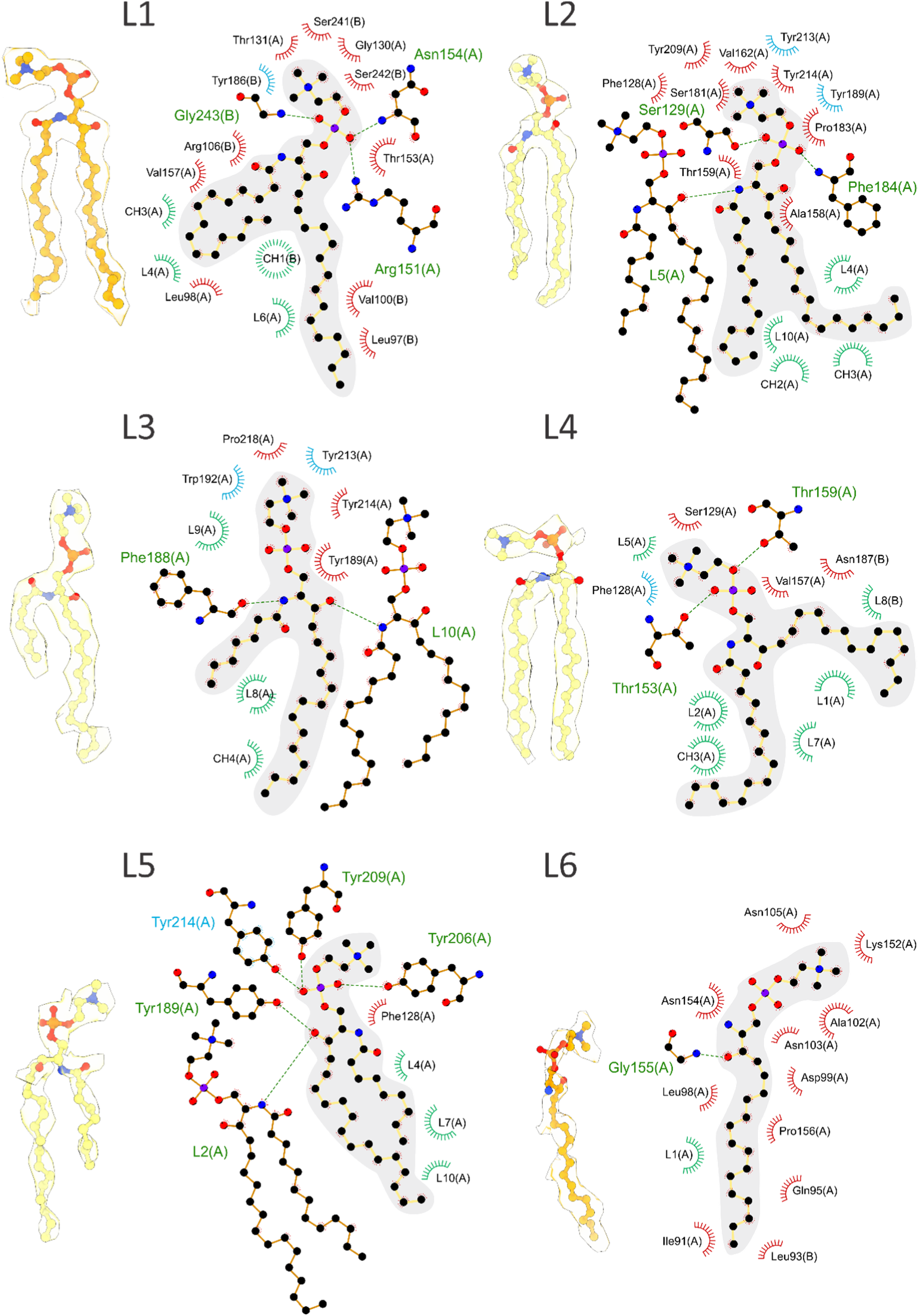
Lipids L1-L6 in the Fav pore structure extracted from large unilamellar vesicles composed of POPG:SM:cholesterol 1:1:1 (mol:mol:mol). Cryo-EM density maps with fitted ball-and-stick models are shown for each lipid (left), with an interaction 2D diagram (right). Hydrogen bonds are represented with green dashed lines. Spoked arcs and circles represent residues (red) and lipids (green) making nonbonded contacts with the lipid. Residues with name in blue or surrounded by blue arcs show cation-π interactions with the choline group. The interaction diagrams were prepared by LigPlot+^55^ and modified by hand after manual inspection.

**Extended Data Fig. 11.**
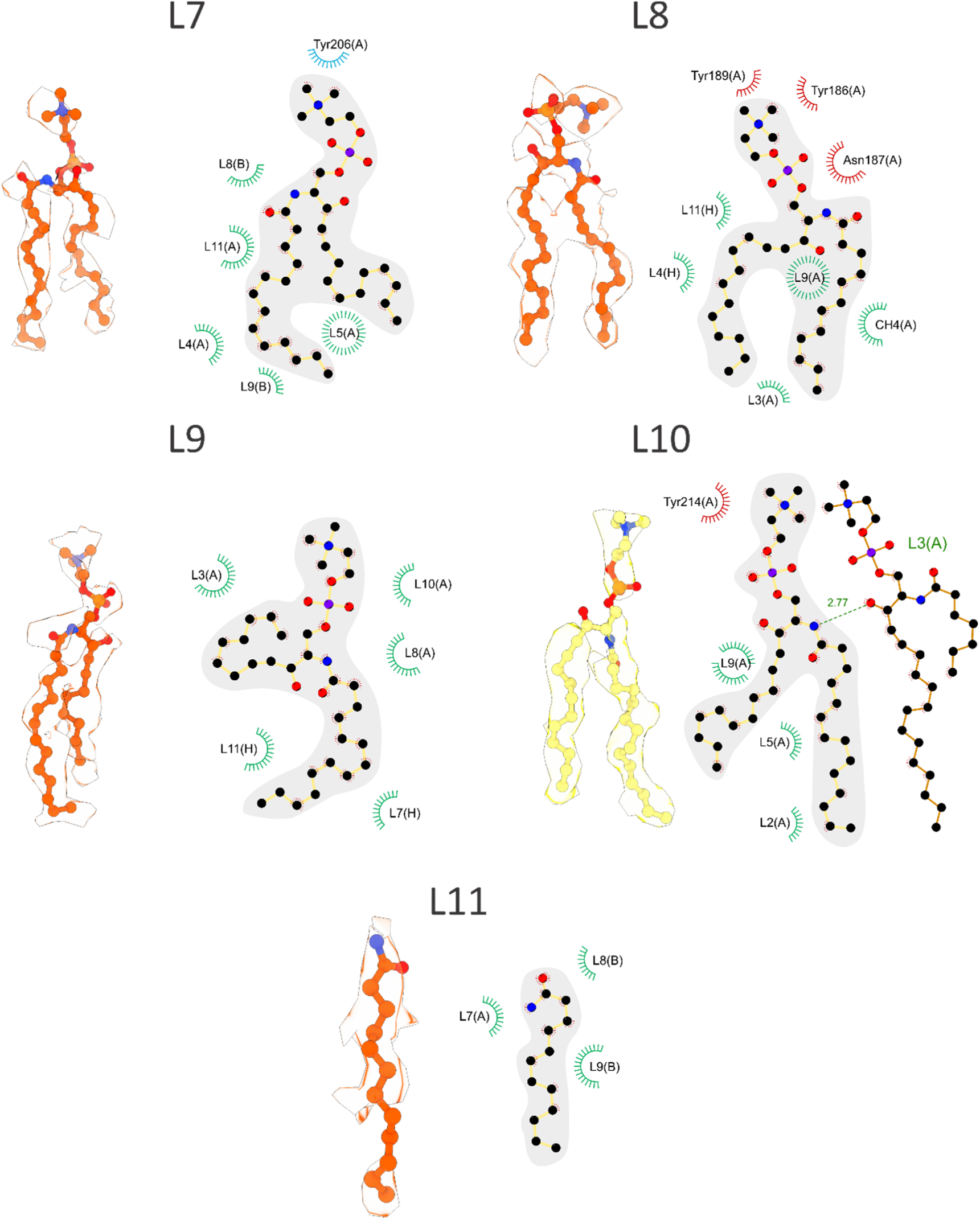
Lipids L7-L11 in the Fav pore structure extracted from large unilamellar vesicles composed of POPG:SM:cholesterol 1:1:1 (mol:mol:mol). Cryo-EM density maps with fitted ball-and-stick models are shown for each lipid position (left), with an interaction diagram (right). Hydrogen bonds are represented with green dashed lines. Spoked arcs and circles represent residues (red) and lipids (green) making nonbonded contacts with the lipid. Residues with name in blue or surrounded by blue arcs show cation-π interactions with the choline group. The interaction diagrams were prepared by LigPlot+^55^ and modified by hand after manual inspection.

**Extended Data Fig. 12.**
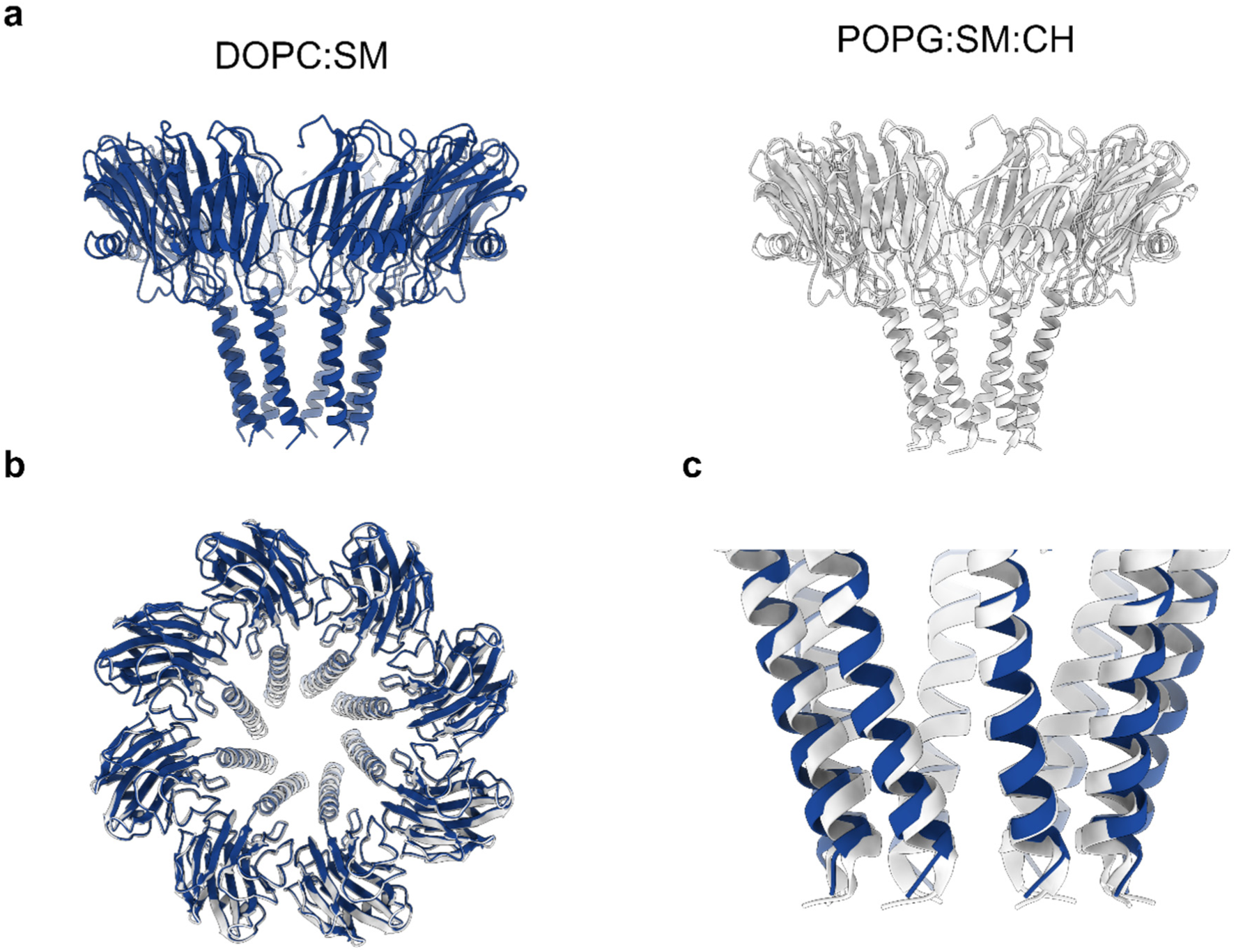
Comparison of pore structures formed in different membranes. Comparison of pores formed on DOPC:SM (grey) and POPG:SM:cholesterol (blue) membranes. **a**, A side-by-side comparison of the two pores. **b**, Top view of aligned pores. **c,** A close-up view at the N-terminal helices of the two pores.

**Extended Data Fig. 13.**
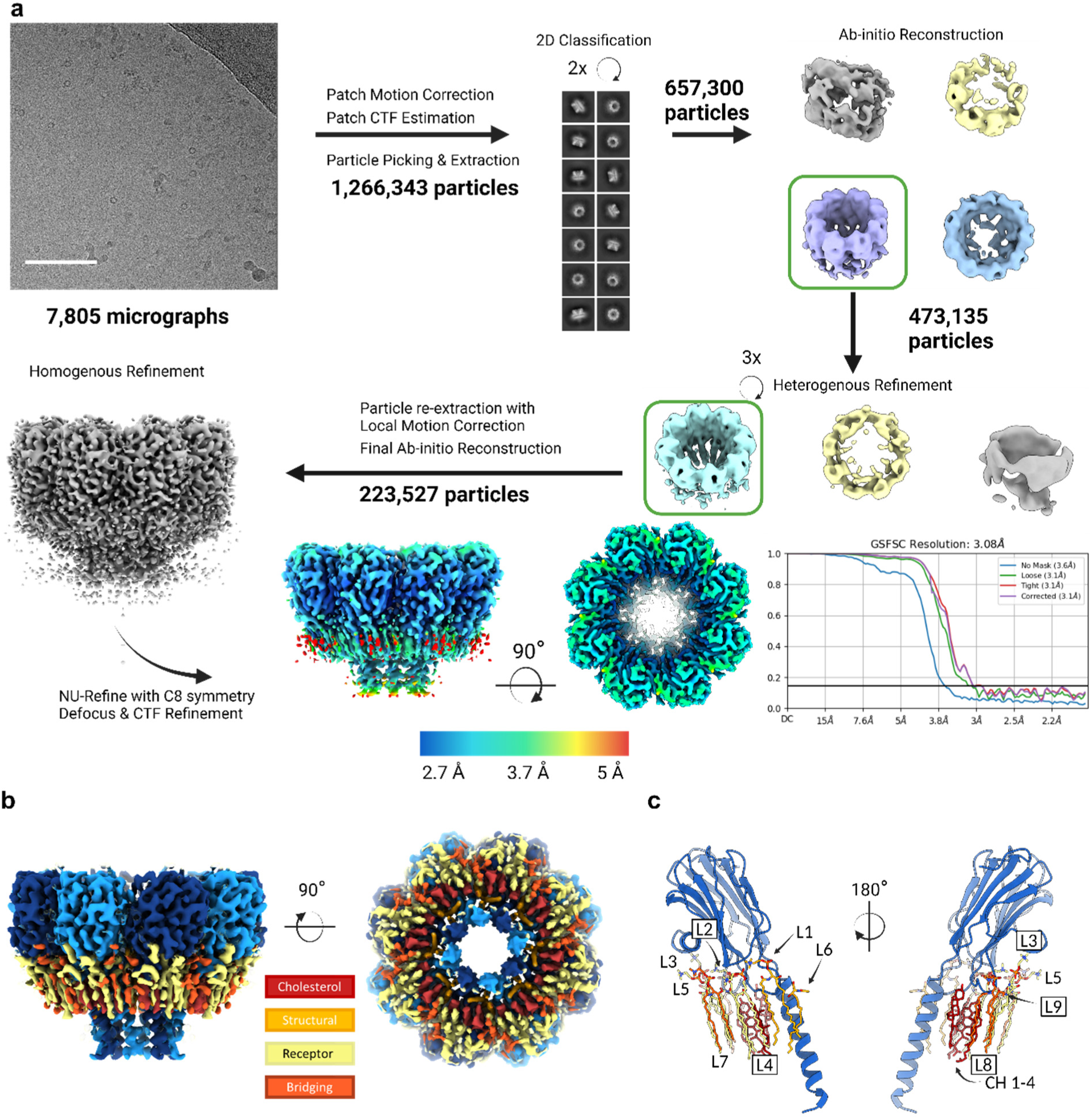
Analysis of wild type Fav pore extracted from large unilamellar vesicles composed of DOPC:SM:cholesterol 1:1:1 (mol:mol:mol). **a**, Cryo-EM data analysis workflow. Details on data analysis are provided in cryo-EM data processing section of Materials and methods. **b,** Cryo-EM map of the Fav pore. Regions corresponding to protomers are coloured blue, while regions corresponding to lipids are coloured by their assigned group. **c**, Cartoon representation of a single protomer with associated lipids at two orientations.

**Extended Data Fig. 14.**
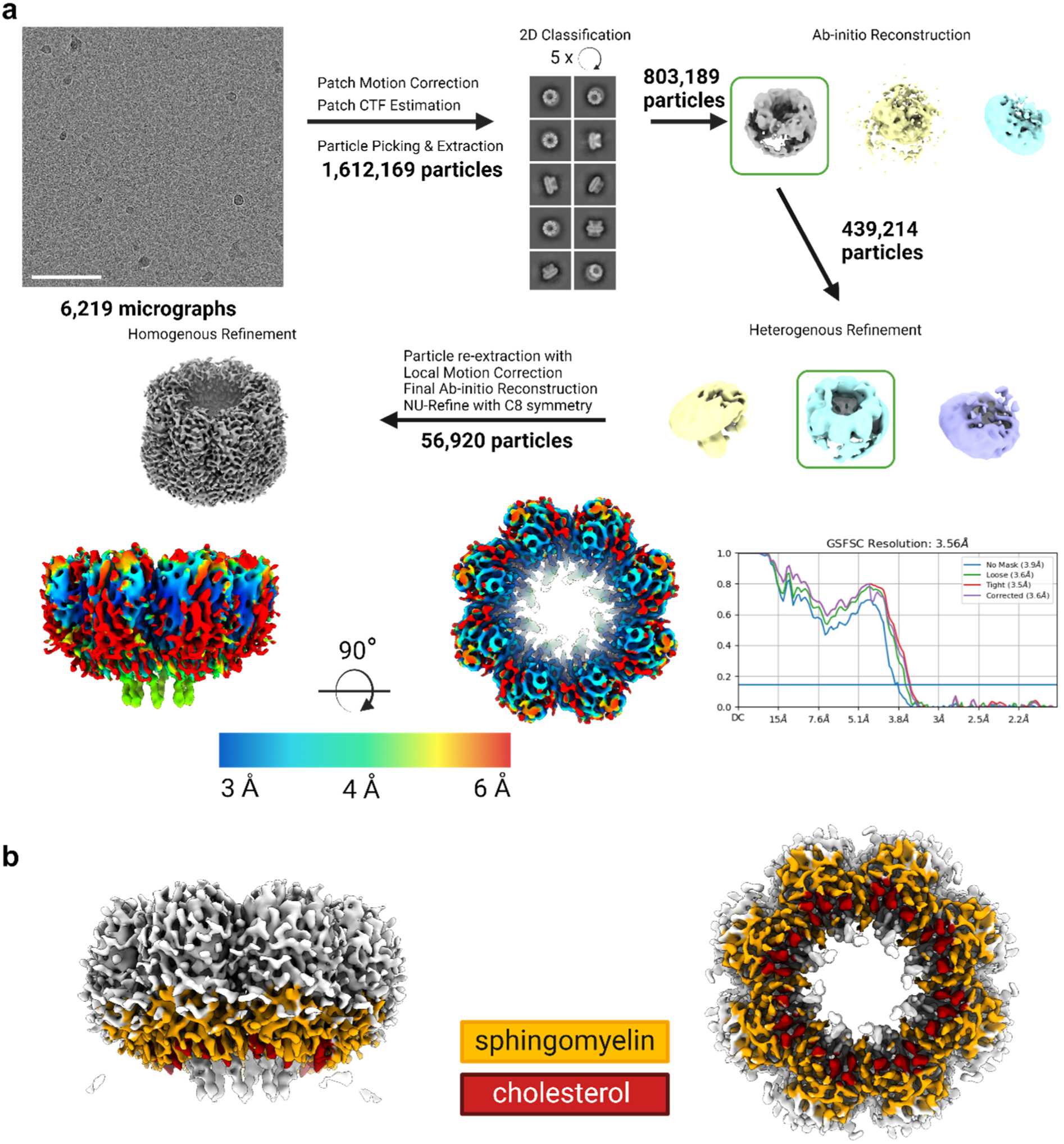
Analysis of wild type Fav pore formed on lipid nanodiscs composed of DOPC:SM:cholesterol 1:1:1 (mol:mol:mol). **a**, Cryo-EM data analysis. Details on data analysis are provided in cryo-EM data processing section of Materials and methods. **b**, Cryo-EM map of Fav pore prepared on nanodiscs. Protein is presented in grey, regions corresponding to cholesterol are colored red and phospholipids are orange.

**Extended Data Fig. 15.**
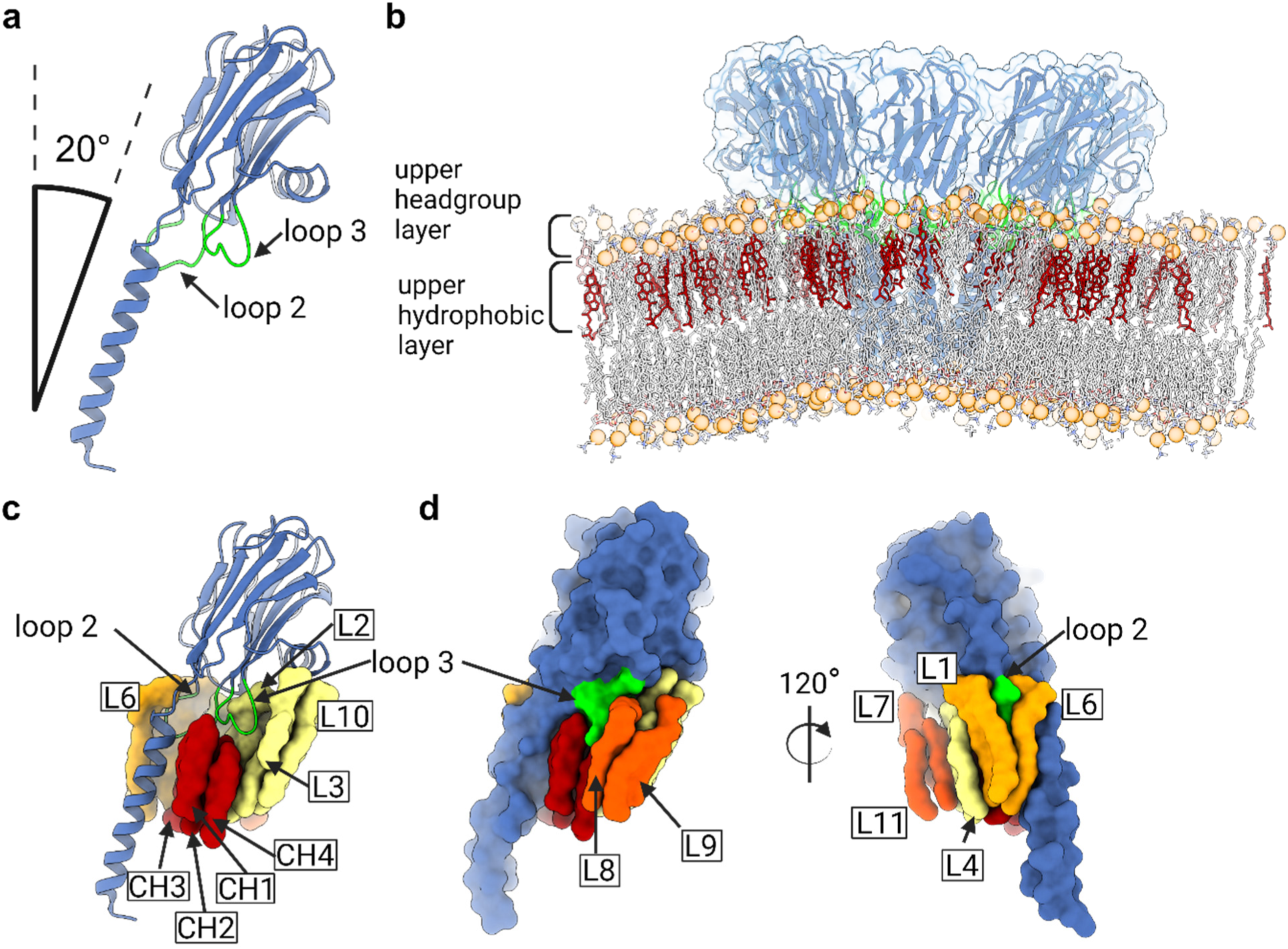
The position of the wild type Fav pore in the membrane. **a**, Fav protomer with membrane binding loops 2 and 3 in green. The angle at which the protomer is placed on the membrane is indicated. **b**, Fav pore (blue) after minimization embedded into a POPC:SM:cholesterol 1:1:1 (mol:mol:mol) membrane that was used for molecular modelling. Loops 2 and 3 of all protomers are shown in green cartoon, orange balls are phosphate groups of lipids and upper leaflet cholesterol molecules are shown in red sticks. **c**, Cartoon representation of a protomer from the pore extracted from large unilamellar vesicles composed POPG:SM:cholesterol 1:1:1 with highlighted loops 2 and 3, with lipids surrounding it. **d**, The surface representation of the Fav protomer with lipids shown at two different angles.

**Extended Data Fig. 16.**
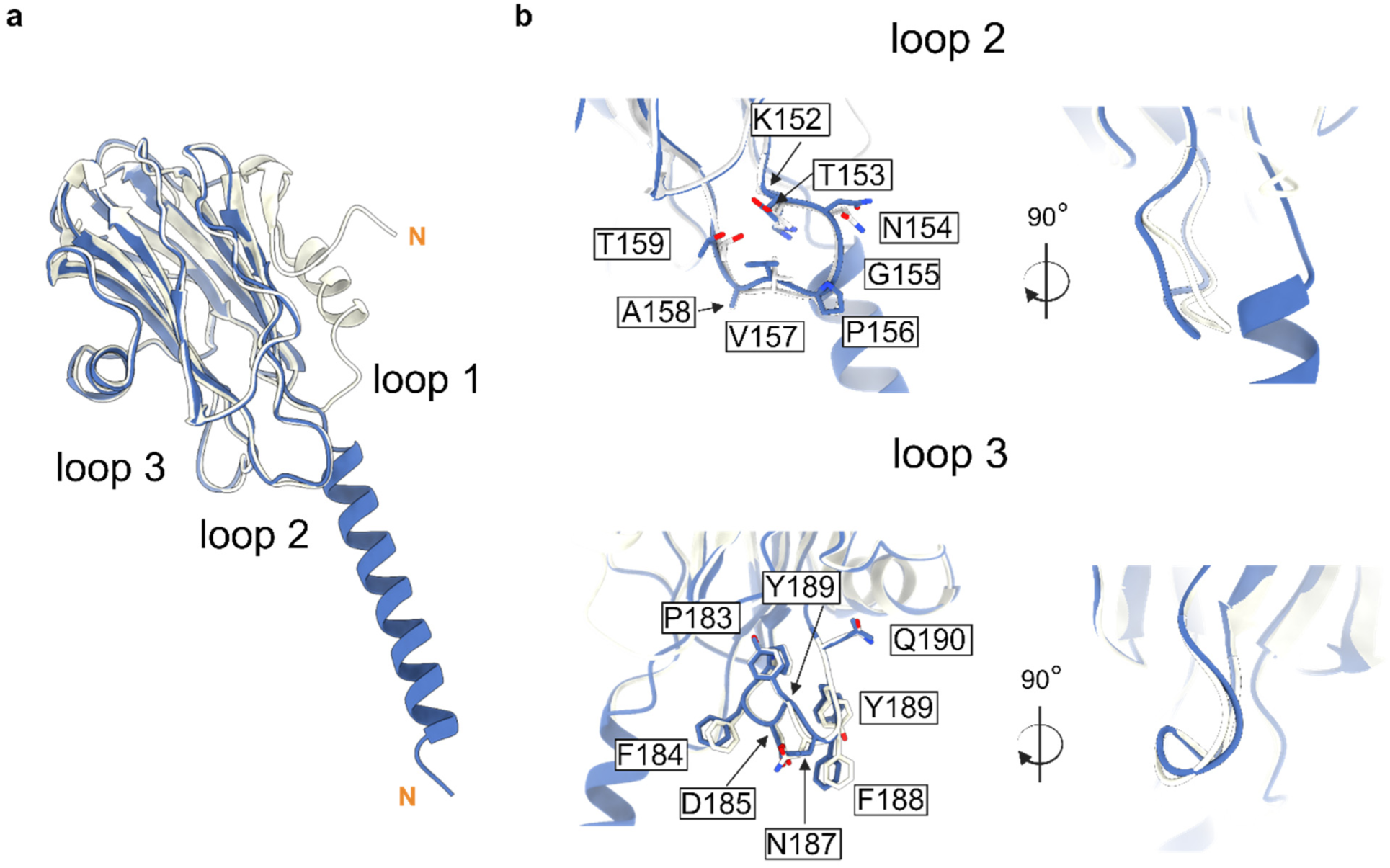
Structural comparison of the soluble Fav monomer and the protomer from the pore. **a**, The overlay of the crystal structure of Δ53Fav monomer (grey cartoon), a single protomer from the pore extracted from large unilamellar vesicles composed POPG:SM:cholesterol 1:1:1 (mol:mol:mol) is shown in blue ribbon. N-terminals are marked with orange N. **b**, Details of loops 2 and 3 are shown with side chains of amino acids presented as sticks and as a cartoon at an 90° angle.

**Extended Data Fig. 17.**
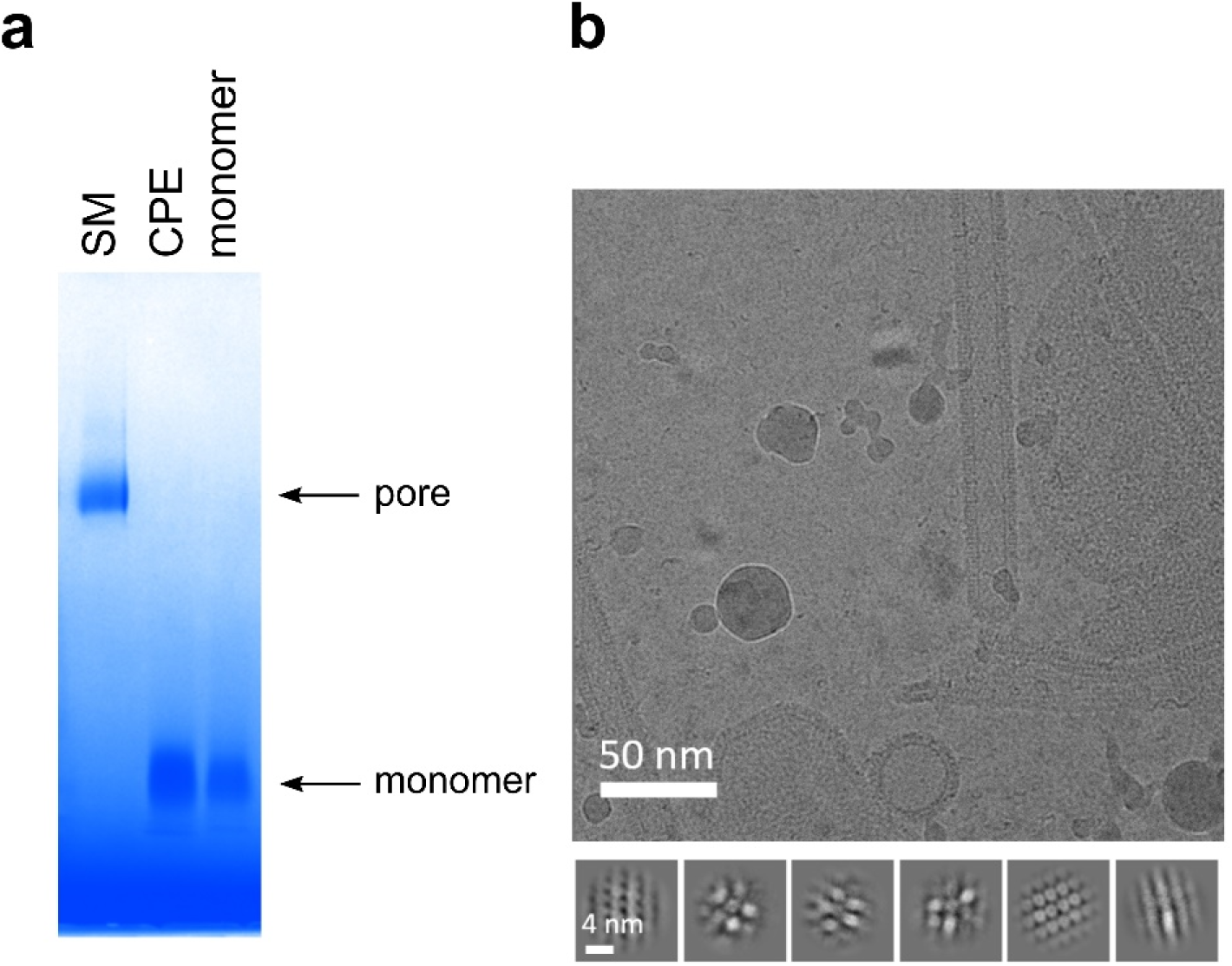
Pore formation by Fav on large unilamellar vesicles containing ceramide phosphoethanolamine (CPE). **a**, Native PAGE of samples obtained by incubating wild type monomeric Fav with large unilamellar vesicles (LUVs) composed of DOPC:SM:cholesterol 1:1:1 (mol:mol:mol) (SM), DOPC:CPE:cholesterol 1:1:1 (mol:mol:mol) (CPE) and without liposomes (monomer), and solubilized by 0.75% lauryl dimethylamine oxide. **b**, Representative cryo-EM micrograph (top) of LUVs composed of DOPC:CPE:cholesterol 1:1:1 after incubation with monomeric wild type Fav. 2D class averages are shown below.

**Extended Data Fig. 18.**
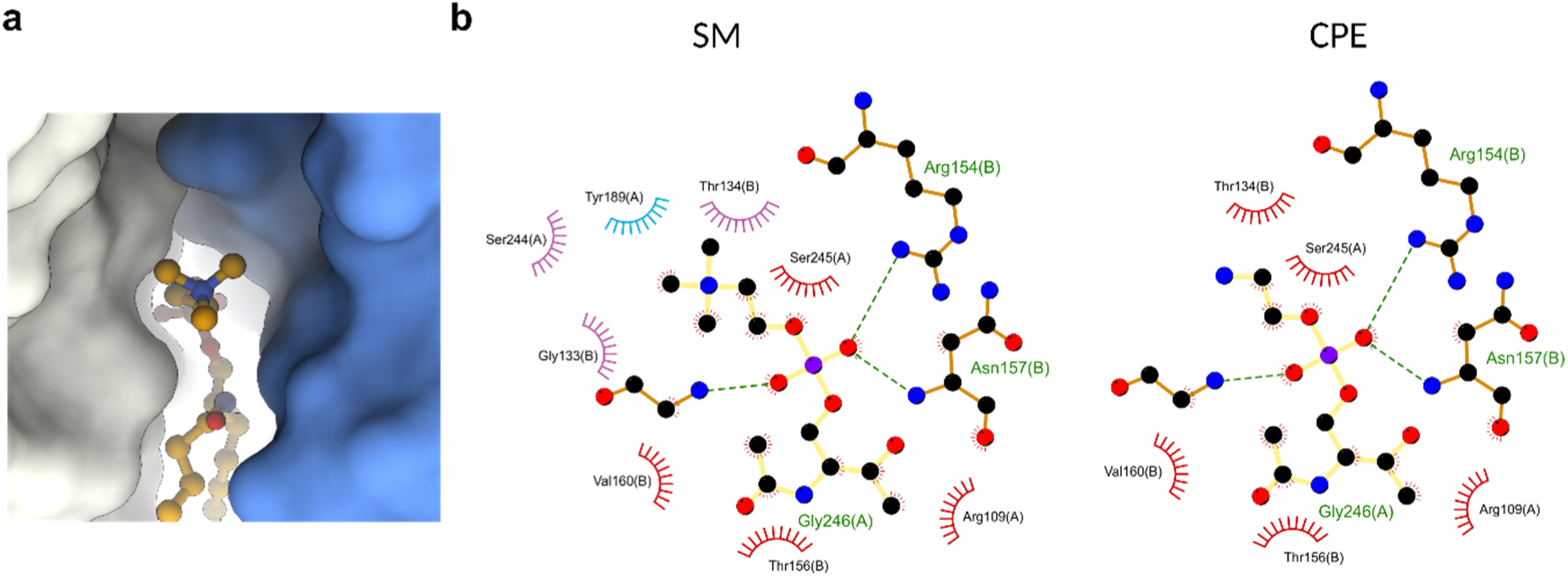
Interactions of the lipid L1 with Fav protomers. **a**, A surface presentation of protomers around headgroup of L1.**b**, Comparison of interaction diagrams of sphingomyelin (SM) at position L1 and potential interaction diagram of ceramide phosphoethanolamine (CPE) at position L1. Spoked arcs and circles represent residues making nonbonded contacts with the lipid. In red are interaction partners present in both lipids, in purple are contacts unique to SM and in blue Y189 forming cation–π interaction with the SM choline.

**Extended Data Fig. 19.**
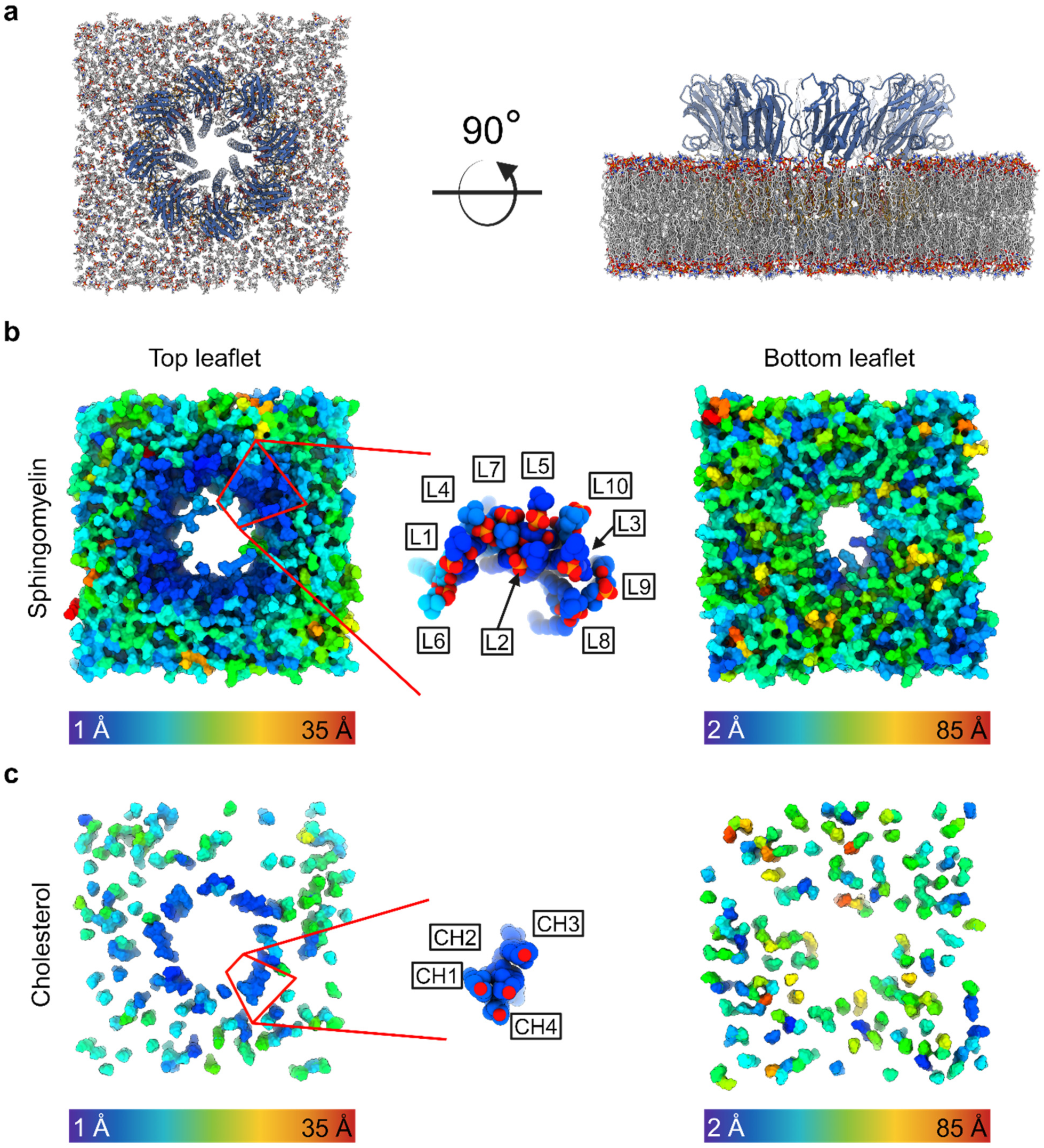
Effect of the pore on lipid mobility. **a**, A look at the inertial frame of molecular dynamics simulation with Fav pore extracted from large unilamellar vesicles composed of POPG:SM:cholesterol 1:1:1 (mol:mol:mol) embedded into POPC:SM:cholesterol 1:1:1 (mol:mol:mol) membrane with 112 lipids at predefined positions (L1-10 and CH1-4). **b** and **c**, Lipids after 1 µs simulation coloured according to the individual lipid displacements are calculated as the average distance travelled by each lipid within 1 µs. Each membrane leaflet is shown independently. **b**, All sphingomyelins with highlighted L1-10 lipids of a single protomer. **c**, All cholesterol molecules with highlighted CH1-4 of a single protomer.

**Extended Data Table 1.** X-ray diffraction data collection and crystallographic refinement statistics for Δ53Fav.

| Data collection | $\Delta 53\text{Fav}$ |
| --- | --- |
| PDB id | 9EYL |
| Resolution range (Å) | 45.8 - 1.5 (1.554 - 1.5) |
| Space group | P 3<sub>2</sub> 2 1 |
| Unit cell dimensions |  |
| a, b, c (Å) | 61.01 61.01 91.82 |
| $\alpha$, $\beta$, $\gamma$ (°) | 90 90 120 |
| Total reflections | 334684 (32351) |
| Unique reflections | 32276 (3196) |
| Multiplicity | 10.4 (10.1) |
| Completeness (%) | 99.93 (99.59) |
| Mean I/sigma (I) | 18.73 (3.25) |
| Wilson B-factor | 11.89 |
| R<sub>merge</sub> (%) | 9.948 (78.93) |
| R<sub>meas</sub> (%) | 10.47 (83.17) |
| R<sub>pim</sub> (%) | 3.232 (26) |
| CC<sub>1/2</sub> | 99.9 (88.7) |
| CC* | 1 (0.969) |
| **Refinement** |  |
| Reflections used in refinement | 32269 (3195) |
| Reflections used for R-free | 1979 (194) |
| Resolution (Å) | 45.8 – 1.5 |
| R-work (%) | 16.33 (23.16) |
| R-free (%) | 18.53 (28.51) |
| CC (work) | 0.969 (0.915) |
| CC (free) | 0.958 (0.880) |
| Number of non-hydrogen atoms | 1773 |
| Protein | 1481 |
| Ligands | 18 |
| Sulfate ion | 10 |
| Tris (114) | 8 |
| water | 274 |
| Protein residues | 184 |
| R. m. s. deviation |  |
| Bond lengths (Å) | 0.008 |
| Bond angles (°) | 0.99 |
| Rotamer outliers (%) | 0.00 |
| Clashscore | 1.00 |
| Average B-factor (Å<sup>2</sup>) | 13.00 |
| macromolecules | 11.23 |
| ligands | 18.04 |
| solvent | 24.68 |

One crystal was used to solve the structure. There was one molecule of monomeric $\Delta 53\text{Fav}$ in the asymmetric unit. Values in parentheses are for the highest-resolution shell.

**Extended Data Table 2.** Interaction surface of Fav protomer from a pore extracted from large unilamellar vesicles composed of POPG:SM:Cholesterol 1:1:1 (mol:mol:mol), and of Fav protomer with removed transmembrane helix with the bound lipids.

| Interaction surface Å<sup>2</sup> |  |  |
| --- | --- | --- |
| Lipid | Whole protomer* | Protomer without transmembrane helix |
| L1 | 325.1 | 229.9 |
| L1(B) | 276.9 | 192.9 |
| L2 | 261.9 | 261.9 |
| L3 | 252.5 | 252.5 |
| L4 | 150.9 | 150.9 |
| L4(B) | 38.0 | 38.0 |
| L5 | 163.7 | 163.7 |
| L6 | 402.9 | 193.9 |
| L6(B) | 93.5 | 0 |
| L7 | 53.9 | 53.9 |
| L8 | 143.6 | 143.6 |
| L9 | 3.9 | 3.9 |
| L10 | 84.9 | 84.9 |
| L11 | 0 | 0 |
| CH1 | 144.4 | 108.9 |
| CH2 | 131.5 | 83.0 |
| CH3 | 152.2 | 69.4 |
| CH4 | 74.7 | 74.7 |
| **Sum of all lipids** | **2759.5** | **2106** |
| **Total available protomer surface** | **9339** | **7443** |

\* The surface was described for the protomer polypeptide chain A. If not marked otherwise, interactions with lipids assigned to bind the protomer A are named. (B) marks the interaction of protomer chain A with the lipid that has been assigned to bind to the protomer chain B.

**Extended Data Table 3.** Cryo-EM data acquisition and refinement statistics.

| Lipid composition | DOPC:SM | DOPC:SM:CH | DOPC:SM:CH<br>nanodiscs | POPG:SM:CH |
| --- | --- | --- | --- | --- |
| **Data Collection** |  |  |  |  |
| Microscope | Krios | Glacios | Glacios | Glacios |
| Magnification | 130,000 | 150,000 | 150,000 | 150,000 |
| Voltage (kV) | 300 | 200 | 200 | 200 |
| Total electron dose (e-/Å<sup>2</sup>) | 40 | 40 | 40 | 40 |
| Defocus range (μm) | -1.0 ~ -2.9 | -0.6 ~ -2.8 | -0.6 ~ -2.6 | -0.6 ~ -2.6 |
| Pixel size (Å) | 1.047 | 0.946 | 0.946 | 0.946 |
| Number of micrographs | 2,140 | 7,805 | 7,533 | 11,470 |
| **Data processing** |  |  |  |  |
| Initial particle number | 802,336 | 1,266,343 | 954,368 | 2,705,265 |
| Final particle number | 174,291 | 223,527 | 193,745 | 118,851 |
| Symmetry imposed | C8 | C8 | C8 | C8 |
| Global map resolution (0.143) (Å) | 2.56 | 2.94 | 3.61 | 2.86 |
| EMDB ID | EMD-50057 | EMD-50060 | EMD-50060 | EMD-50059 |
| **Model Building** |  |  |  |  |
| Associated PDB ID | 9EYM | 9EYN |  | 9EYO |
| FSCmodel-to-map (0.5) (Å) | 2.99 | 3.30 |  | 3.28 |
| Composition |  |  |  |  |
| Protein chains | 8 | 8 |  | 8 |
| Atoms | 13664 | 15296 |  | 16160 |
| Protein residues | 1448 | 1448 |  | 1456 |
| Ligands* | 48 (A1H8M) | 32 (CLR),<br>72 (A1H8M) |  | 32 (CLR),<br>90 (A1H8M) |
| Bonds (R.M.S.D.) |  |  |  |  |
| Length (Å) | 0.012 (16) | 0.012 (16) |  | 0.012(0) |
| Angles (°) | 1.830 (24) | 1.836 (48) |  | 1.884 (48) |
| B-factors (Å<sup>2</sup>) (min/max/mean) |  |  |  |  |
| Protein residues | 77.29/226.90/124.57 | 136.80/238.01/179.18 |  | 73.19/195.42/101.35 |
| Ramachandran plot (%) |  |  |  |  |
| Outliers | 0.00 | 0.00 |  | 0.00 |
| Allowed | 1.68 | 1.68 |  | 1.67 |
| Favored | 98.32 | 98.32 |  | 98.33 |
| Rotamer outliers (%) | 0.68 | 0.68 |  | 2.72 |
| Cβ outliers (%) | 0.00 | 0.00 |  | 0.00 |
| MolProbity score | 1.01 | 0.89 |  | 0.97 |
| Clash Score | 2.30 | 1.47 |  | 0.73 |

\*All phospholipids were assigned as sphingomyelin (A1H8M in the PDB file for simplicity). During the study, we were able to show that L1-L5 are indeed sphingomyelin. CLR: three letter abbreviation in the PDB file for cholesterol.

